# Extracellular ATP is an environmental cue in bacteria

**DOI:** 10.1101/2024.01.24.577065

**Authors:** Sophie Tronnet, Vikash Pandey, Miriam Lloret-Berrocal, Mario Pérez-del-Pozo, Niklas Söderholm, Carlos Hernández-Ortego, Oliver Billker, Anders Nordström, Andrea Puhar

**Affiliations:** The Laboratory for Molecular Infection Medicine Sweden (MIMS), 901 87 Umeå, Sweden; Umeå Centre for Microbial Research (UCMR), 901 87 Umeå, Sweden; Department of Molecular Biology, Umeå University, 901 87 Umeå, Sweden; Department of Chemistry, Umeå University, 901 87 Umeå, Sweden; Umeå Plant Science Centre, 901 87 Umeå, Sweden; Swedish Metabolomics Centre, 907 36 Umeå, Sweden; Wellcome-Wolfson Institute for Experimental Medicine, School of Medicine, Dentistry and Biomedical Sciences, Queen’s University Belfast, BT9 7BL Belfast, United Kingdom

## Abstract

In animals and plants extracellular ATP (eATP) functions as signal and regulates the immune response. During inflammation intestinal bacteria are exposed to elevated eATP originating from the mucosa. However, whether bacteria respond to eATP is unclear. Here we show that non-pathogenic *Escherichia coli* responds to eATP by modifying its transcriptional and metabolic landscapes. A genome-scale promoter library showed that the response is time-, concentration-, and medium-dependent and ATP-specific. The second messenger cAMP and genes related to metabolism, quorum sensing, and envelope stress were regulated downstream of eATP. Metabolomics confirmed that eATP triggers enrichment of compounds with bioactive properties on the host or bacteria. Combined genome-scale modelling revealed global metabolic and biomass building blocks modifications. Consequently, eATP altered the sensitivity to antibiotics and antimicrobial peptides. Finally, in pathogens eATP controlled virulence factor expression. Our results indicate that eATP is an environmental cue in prokaryotes which broadly regulates physiology, antimicrobial resistance, and virulence.

## Introduction

Extracellular ATP (eATP) has emerged as a key signal in lower and higher eukaryotes, including protozoa, animals, plants, and fungi^1–6^. In animals and plants eATP regulates physiological responses^1,2^. Additionally, eATP controls damage, stress or inflammatory responses in protozoa, animals, plants and fungi upon tissue injury or alteration^1–5,7,8^. In contrast, the existence of purinergic signalling in the earlier forms of life, Bacteria and Archaea, remains largely unexplored^1^.

Only a handful of studies addressed the effect of eATP on bacteria at the phenotypic level. In *Streptomyces* exogenous ATP promoted differentiation, possibly via cytosolic ATP-consuming enzymes^9–11^. Other studies investigated the role of added ATP in biofilms and/or growth in several organisms, partially with opposing results. Effects on biofilms were linked to release of biofilm-composing DNA upon eATP-induced lysis^12^, inhibition of twitching motility by reducing surface-assembled Type IV pili via eATP hydrolysis^13^, chelation of biofilm-composing cations by eATP^14^, or not further dissected^15^. Effects on growth were not explained^12,16,17,18^, except for eATP-dependent iron deficiency in one instance^19^. Finally, ATP supplementation increased survival of non-growing bacteria^20^. Overall, whether eATP acts as environmental cue to which bacteria respond remains unsolved.

Bacteria are typically exposed to low concentrations of eATP, partly originating from themselves. In fact, eATP is present in the nano- to micromolar range in culture supernatants^21^ or homogenates^22^ of murine faeces and many isolated bacteria in culture^13,15,18,20,22–28^ at steady state. Additionally, eATP is found in bacterial supernatants upon osmotic shock^29^, treatment with antimicrobial peptides^30,31^ or antibiotics^18,28,32–34^, or compromised outer membrane stability^35^. Likewise, during homeostasis eATP is found in the gut lumen of mammals at low concentration^17,34,36–41^.

Importantly, intestinal eATP dramatically increases during bacterial, viral, or parasitic infection, protist colonisation, and inflammation. This implies that under these conditions, commensal and pathogenic microbes are exposed to heightened concentrations of eATP in the gut lumen. Indeed, we recently discovered that enteropathogenic bacteria trigger connexin-dependent secretion of ATP from the intestinal epithelium, *via* activation of the mechanosensor PIEZO1, as an early alert response, which induces potent gut inflammation^36,42^. Rotavirus also initiates epithelial purinergic signalling^43^. During Shiga toxin-producing enterohaemorrhagic *E. coli*^44^ or *Citrobacter rodentium* infection^22^ and *Tritrichomonas musculis* colonisation^45^ high eATP is present in faeces. Helminths induce ATP release in the gut^46^. Further, intestinal eATP rises during experimental inflammatory bowels diseases (IBD)^37,47–49^ and after irradiation in graft-versus-host-disease^50^ and small intestinal fibrosis^51^. Ultimately, any process causing cell damage results in ATP release^1^. Taken together, these drastic changes in intestinal eATP concentration may serve as environmental signal to bacteria.

Here we show that eATP regulates the transcriptional and metabolic landscapes of non-pathogenic *E. coli* at physiologically relevant concentrations. The response to eATP presents the characteristic of bacterial transcriptional and metabolic networks, and modulates the production of bioactive compounds, biofilm formation, and susceptibility to antimicrobials. Furthermore, several gut pathogens and pathobionts respond to eATP by regulating expression of virulence or fitness factors. Together, our results indicate that eATP is an environmental cue that globally modifies the physiology of harmless and pathogenic bacteria.

## Results

### eATP triggers a robust transcriptional response in non-pathogenic *E. coli*

To investigate the ability of bacteria to sense and respond to eATP, we tested whether eATP induces a transcriptional response using a plasmid-based fluorescent reporter library in non-pathogenic *E. coli* K-12 MG1655^52^. This collection can unravel the dynamic activity of approximately 2000 promoters (75% of *E. coli* promoters), allowing accurate quantification of gene expression induction or repression seen as GFP fluorescence normalised to the number of bacteria.

We screened this library for a response to eATP in rich (LB) and minimal medium (M9) upon treatment for 3 h with 1 mM ATP, a concentration in the range of inflammatory environments (Figure S1A). Addition of ATP led to robust up- or down-regulation of promoters in both media, with more modulated in M9 than in LB (Figure 1A-D, examples in Figure S1B-G, full list of genes and fold changes in Table S1). We identified 291 and 476 modulated promoters in LB and M9, respectively (Figure 1E), with 121 promoters modulated by eATP in both media, of which 46 genes were upregulated and 27 downregulated (Figure 1F, G). These screens were highly reproducible (Figure S1H), and addition of ATP neither altered bacterial growth (Figure S1I-L) nor the pH of the medium (Figure S1M).

**Figure 1.**
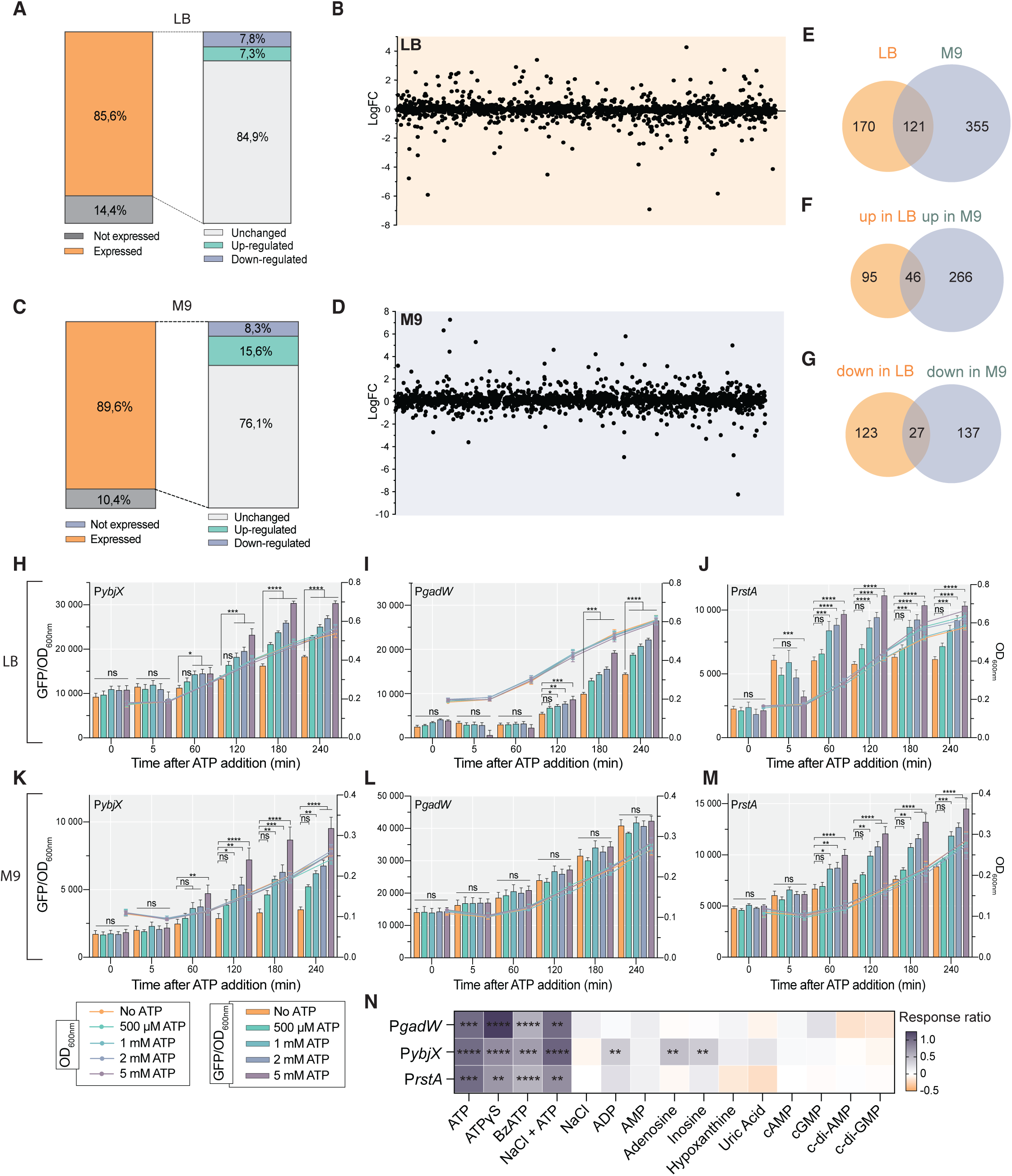
eATP triggers a strong medium-, dose-, and time-dependent transcriptional response in non-pathogenic *E. coli*. **A** and **C**, Number of eATP-regulated promoters in percentage with respect to unstimulated controls from fluorescence reporter screens (n=2) performed in LB or M9 in *E. coli* MG1655, 180 min after addition of 1 mM ATP. **B** and **D**, Log fold change of promoter activity upon exposure to 1 mM eATP 180 min after ATP addition in LB or M9 in *E. coli* MG1655, shown as mean of independent experiments (nζ2). **E**-**G**, Comparison of the number of (**E**) all differentially expressed promoters in *E. coli* MG1655, (**F**) all upregulated promoters, and (**G**) all down-regulated promoters 180 min after addition of 1 mM ATP in LB or M9. **H**-**M**, Promoter activity of selected upregulated promoters P*ybjX*, P*gadW* and P*rstA* controlling the expression of the reporter GFP in the presence of increasing concentrations of added ATP in *E. coli* MG1655. Strains were grown in LB (upper panels) or M9 (lower panels). Shown is the ratio of GFP fluorescence to the number of bacteria determined by absorbance at 600 nm over time (columns representing the mean with s.e.m., left Y axes), as well as the bacterial growth (continuous lines representing means with s.e.m., right Y axes). Statistical analysis: two-way ANOVA with Dunnett’s multiple comparison test; *P < 0.05, **P < 0.01, ***P < 0.001 and ****P < 0.0001; ns, not significant. **N**, Activity of selected upregulated promoters as in H-M in the presence of eATP-related compounds at 1 mM in *E. coli* MG1655 in LB. The expression ratio was calculated comparing the fluorescence/OD_600nm_ in LB to fluorescence/OD_600nm_ in LB + compound. The response ratio of ATP is set to 1. Shown is the mean with s.e.m. of ≥ 3 independent experiments performed in duplicates or triplicates. Statistical analysis: two-way ANOVA with Dunnett’s multiple comparison test with respect to unstimulated control; **P < 0.01, ***P < 0.001 and ****P < 0.0001; ns, not significant (omitted).

We confirmed the activity of two promoters upregulated in LB in their native context by constructing chromosomal fusions with superfolder GFP. Both endogenous promoters of *gadW* and *cbpA* (controlling expression of a DNA-binding transcriptional dual regulator and a DnaK co-chaperone) were significantly upregulated (fold change::2) 5 and 30 min after eATP addition, respectively (Figure S1N-P).

To investigate whether the transcriptional response to eATP in *E. coli* MG1655 is dose- and time-dependent, we selected three reporters that were significantly upregulated by eATP at 3 h in LB and/or M9: P*ybjX*, controlling the expression of a DUF535 domain-containing protein; P*gadW*; and P*rstA*, controlling the DNA-binding transcriptional regulator of the two-component system RstAB. We applied increasing concentrations of eATP and measured the promoter activity in LB (Figure 1H-J) and M9 (Figure 1K-M) over time. This showed that the response to eATP was dose-dependent for all tested promoters in both media after 60 min of stimulation or longer, except for P*gadW*, which was not eATP-regulated in M9 (Figure 1L). These tests showed that transcriptional response to eATP is dynamically regulated in the high micromolar range.

To determine whether the transcriptional response is specific to ATP, we challenged *E. coli* MG1655 in LB with a panel of ATP-related compounds at 1 mM (Figure 1N). Because eATP is unstable, we tested the non-hydrolysable analogue ATPγS. This showed an induction of P*gadW*, P*somA*, or P*rstA* similar to ATP compared to the medium. BzATP, a nonselective agonist of mammalian P2X purinergic receptors (which sense ATP)^8,53^, also triggered a transcriptional response comparable to ATP. We controlled for the introduction of sodium from the ATP stock by adding sodium chloride alone or together with ATP, which showed no effect. Moreover, we tested the ATP degradation products ADP, AMP, adenosine, inosine, hypoxanthine, and uric acid^8^, which overall failed to regulate P*gadW*, P*somA*, or P*rstA*. Finally, supplementation of nucleotides with known intracellular signalling function in bacteria, the second messengers cAMP, cGMP, c-di-AMP, and c-di-GMP^54^, did not activate a transcriptional response. Overall, these experiments indicate that intact eATP induces transcriptional activation from the cell surface, because alike mammalian cells *E. coli* does not internalise ATP^20,55^.

In summary, these results show that the intestinal non-pathogenic bacterium *E. coli* specifically responds to eATP at physiologically relevant concentrations by modifying its transcriptional landscape in a medium-, dose- and time-dependent manner.

### eATP elicits cAMP production and regulates the physiology of non-pathogenic *E. coli*

We used the data from the transcriptional reporter library screen to determine processes regulated by eATP in *E. coli* MG1655. We noticed that among the eATP-induced promoters, 36 in LB and 59 in M9 are regulated by the cAMP receptor protein (CRP) (Figure 2A). Therefore, we tested the the cAMP concentration by ELISA that and found significant cAMP increases upon eATP addition in LB and decreases in M9 (Figure 2B-C), suggesting that eATP triggers an intracellular signalling cascade. Indeed, the second messenger cAMP is produced downstream of environmental signals and regulates carbon metabolism, stress response pathways, virulence, and biofilm formation^56^. We chose to test potential cAMP effects by quantifying biofilm formation using crystal violet, which decreased in LB and increased in M9 when treated with eATP (Figure 2D-E).

**Figure 2.**
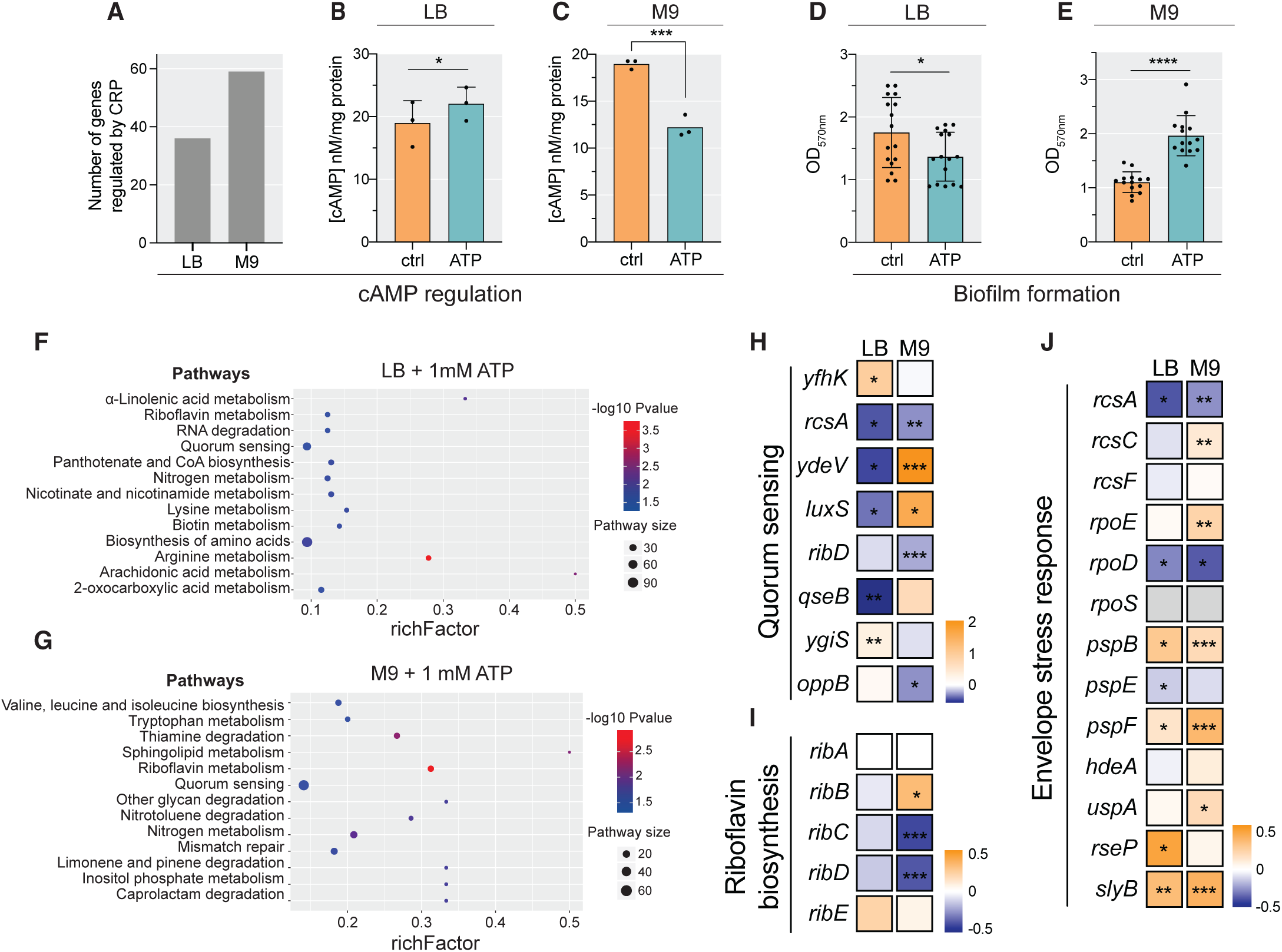
Pathway analysis of the eATP-triggered transcriptional response in non-pathogenic *E. coli*. **A**, Count of modulated CRP-regulated genes in *E. coli* MG1655 stimulated with 1 mM eATP for 180 min in LB or M9, from data in Table S1. **B-C** , Determination of the intracellular cAMP concentration by ELISA normalised to the number of bacteria seen as protein concentration in lysates from *E. coli* MG1655 stimulated with 1 mM eATP for 180 min in LB or M9. Statistical analysis: Student’s *t* test; *P < 0.05 and ***P < 0.001. **D**-**E**, Biofilm formation assays using crystal violet staining in *E. coli* MG1655 stimulated with 1 mM eATP before 24 h static growth in LB or M9. Statistical analysis: Student’s *t* test; *P < 0.05 and ****P < 0.0001. **F**-**G**, KEGG pathway enrichment analysis on the up- and downregulated genes in LB or M9 180 min after 1 mM ATP addition in comparison to unstimulated bacteria, from data in Table S1. **H**, Analysis of quorum sensing-related genes in *E. coli* MG1655 stimulated with 1 mM eATP for 180 min in LB or M9, from data in Table S1. Statistical analysis: two-way ANOVA with Dunnett’s multiple comparison test; *P < 0.05, **P < 0.01 and ***P < 0.001; ns, not significant (omitted). **I**, Analysis of genes related to riboflavin biosynthesis in *E. coli* MG1655 stimulated with 1 mM eATP for 180 min in LB or M9, from data in Table S1. Statistical analysis: two-way ANOVA with Dunnett’s multiple comparison test; *P < 0.05 and ***P < 0.001; ns, not significant (omitted). **J**, Analysis of genes related to the envelope stress response biosynthesis in *E. coli* MG1655 stimulated with 1 mM eATP for 180 min in LB or M9, from data in Table S1. Statistical analysis: two-way ANOVA with Dunnett’s multiple comparison test; *P < 0.05, **P < 0.01, ***P < 0.001 and ****P < 0.0001; ns, not significant (omitted).

To gain a deeper understanding of the response to eATP in non-pathogenic *E. coli* MG1655, we analysed the function of eATP-modulated genes by pathway prediction. Several regulated genes encode unknown or poorly characterized proteins (Table S1). We analysed the differentially expressed genes (DEGs) using Panther^57^ (Figure S1Q, Table S1). The modulated DEG were found in the subcategories: binding, catalytic activity, cellular processes, metabolic process, cellular anatomical entity, metabolite interconversion enzyme and, to a lesser extent, transporter and gene-specific transcriptional regulator, showing that eATP impacts bacterial gene regulation, metabolism, and composition.

KEGG analysis revealed 13 and 15 significantly enriched pathways in LB and M9, respectively, including the metabolism of fatty acids, vitamins, co-factors and proteinogenic amino acids (Figure 2F-G, Table S1). The metabolism of different types of vitamins, amino acids and fatty acids was predicted in LB and M9. By contrast, quorum sensing, riboflavin and nitrogen metabolism were enriched in both media. Therefore, we looked at promoters involved in quorum sensing. As expected from the role of quorum sensing in biofilm formation (Figure 2D-E), eATP oppositely regulated quorum sensing genes in LB and M9 (Figure 2H). Likewise, as an example of predicted eATP-modulated vitamin metabolism pathways, we found that the riboflavin biosynthesis-associated promoters P*ribC* and P*ribD* were significantly downregulated in M9 (Figure 2I). Finally, we noticed that eATP modulated promoters related to the envelope stress response^58^ in both media (Figure 2J). Specifically, eATP regulated the Rcs and Psp systems, which detect lipopolysaccharide, peptidoglycan or lipoprotein defects and damage to the inner membrane, respectively^58^. Similarly, in LB and M9, eATP significantly activated P*slyB*, involved in cell-envelope proteostasis and integrity^59^.

Together, these experiments show that eATP induces production of a secondary messenger and genes important to metabolite production, biofilm formation, and envelope stress.

### eATP triggers a strong metabolic response in non-pathogenic *E. coli*

Our data indicated that the gene expression programmes induced by eATP have a strong impact on bacterial physiology and metabolism. To validate this finding, we assessed the changes in intracellular (pellet) and extracellular (supernatant) metabolites of bacteria grown in LB or M9 upon exposure to eATP by untargeted metabolomics using liquid chromatography‒mass spectrometry (LC‒MS/MS). We confirmed the identity of shown compounds using an analytical standard library. We only measured extracellular metabolites in M9 because LB cannot be employed in LC-MS/MS.

In unstimulated *E. coli* MG1655, we detected 57 and 53 intracellular metabolites in LB and M9, respectively, with 51 present in both media. Further, we identified 89 extracellular metabolites, with 61 only present in the supernatant but not the pellets, while 23 metabolites were found in all conditions (Figure 3A and Table S2). We then compared the metabolome of bacteria stimulated with 1 mM ATP for 3 h with time-matched unstimulated controls. Partial least square-discriminant analysis (PLS-DA) distinguished untreated and eATP-treated *E. coli* (Figure 3B-D), indicating a clear effect of eATP on the metabolome. Univariate analysis of these data revealed that eATP triggers a potent shift in the metabolic landscape (Figure 3E-J, Table S2), with 5–16% of detected metabolites significantly modulated (≥1.5 FC, *P* ≤ 0.05) in response to eATP. The metabolic changes upon ATP addition in M9 were more substantial than in LB with 9/53 and 4/57 significantly modulated metabolites in pellets, respectively (Figure S2A-D, Table S2), confirming the medium-dependent effect of eATP we observed in transcriptional assays (Figures 1-2). We found 11/89 significantly eATP-modulated metabolites in the supernatant (Figure S2E-F, Table S2).

**Figure 3.**
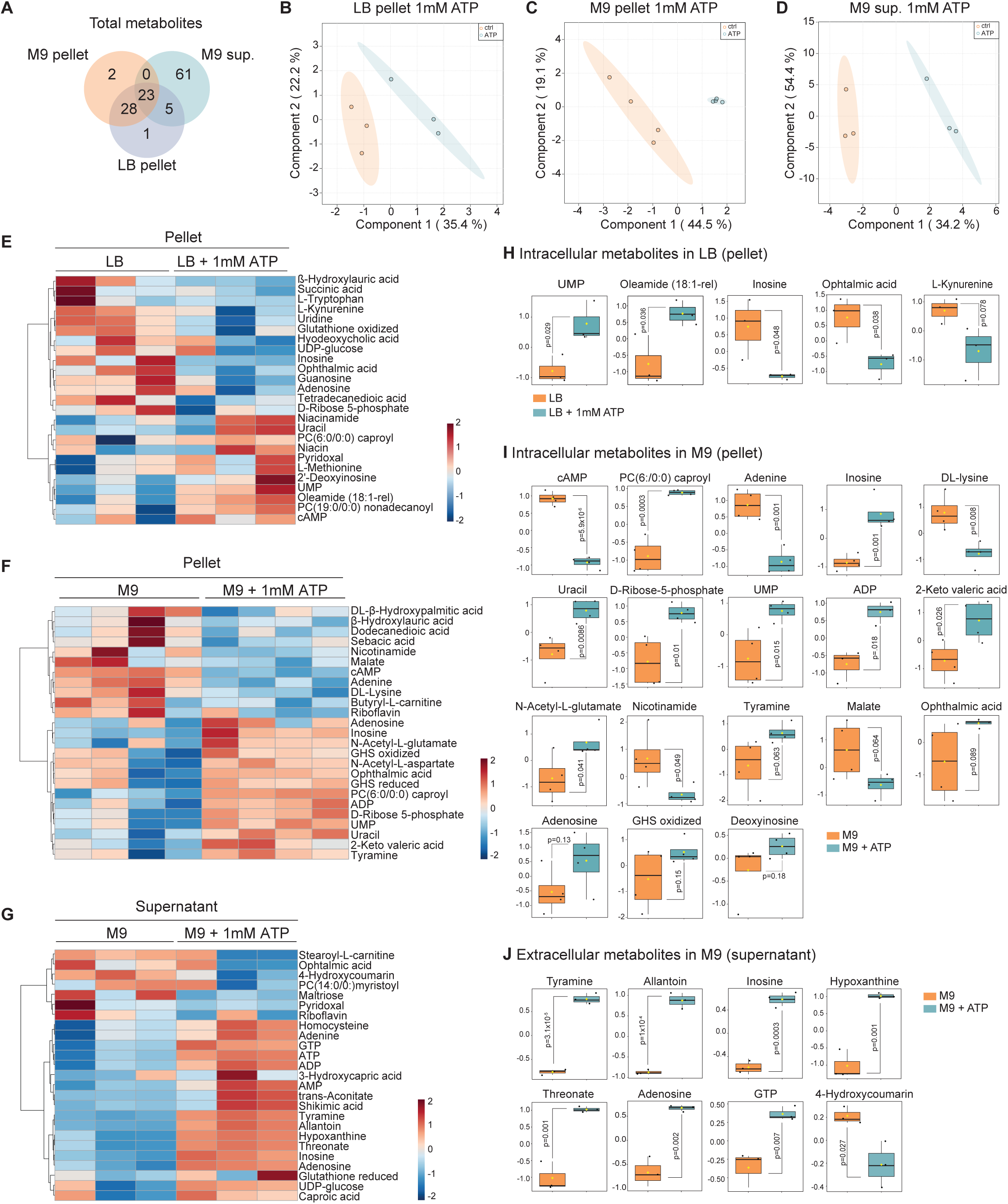
eATP triggers a strong metabolic response in non-pathogenic *E. coli*. **A**, Comparison of the total number of common or unique detected metabolites from *E. coli* MG1655 grown in LB or M9 after 180 min in the presence of 1 mM added ATP. LB pellet: n=3 and 57 metabolites, M9 pellet: n=4 and 53 metabolites, M9 supernatant: n=3 and 89 metabolites. **B-D** , Partial Least Squares discriminant analysis (PLS-DA) scores plots of components one and two, comparing metabolomic samples from *E. coli* MG1655 treated with 1 mM ATP for 180 min versus the unstimulated control in (**B**) the pellet in LB (n=3), (**C**) the pellet in M9 (n=4), and (**D**) the supernatant in M9 (n=3). **E**, Comparison of fold changes in abundance of top 25 metabolites based on PLS-DA analysis from samples in (**B**). **F**, Comparison of changes in abundance of top 25 metabolites based on PLS-DA analysis from samples in (**C**). **G**, Comparison of fold changes in abundance of top 25 metabolites based on PLS-DA analysis from samples in (**D**). **H**-**J**, Changes of relative concentrations of selected modulated intracellular metabolites from samples in (**B** and **C**) and extracellular metabolites from samples in (**D**). The boxplots show the normalised values (mean and s.d.). The Y-axis represents the normalised peak area of metabolites. Black dots represent the normalised concentrations of all samples of each metabolite. The notch indicates the 95% confidence interval around the median. The mean concentration of each group is shown by a yellow diamond. Statistical analysis: two-sided Welch’s t test; raw P-values are indicated in each panel.

Overall, the most recurrently modulated metabolites were intermediates of fatty acid, co-factor, small organic acid, pentose phosphate, vitamin, nucleotide and nucleoside, proteinogenic and non-proteinogenic amino acid biosynthesis, and redox regulation (Figure 3E-J, Table S2), in keeping with the predicted regulation of metabolic pathways based on eATP-dependent transcriptional changes (Figure 2F-G). To identify the most characteristic metabolites in LB and M9, we calculated VIP scores, ranking the metabolites according to their influence on the separation of the treatment groups (Figure S2B, S2D, S2F). To analyse these differences in depth, we constructed metabolic models (see below). Among the metabolites not integrated in the metabolic model, ophthalmic acid was significantly decreased in LB (Figure 3H, Table S2). In M9, cAMP (as seen in Figure 2C) and lysine were strongly decreased, while the membrane lipid PC-caproyl was upregulated by eATP (Figure 3I, Table S2). Further, the analysis showed that the upregulated biogenic amine tyramine was the top ranked metabolite in the VIP analysis in the M9 supernatant (Figure 3J, Figure S2E-F). Of note, allantoin, AMP, threonate and GTP also accumulated in the supernatant. Many of these compounds have potential effects on the host or other gut microbes (see Table S2 and Discussion) and may be produced in eATP-high inflammatory environments.

To observe metabolite changes at low eATP, alike the dose-dependent transcriptional response (Figure 1H-M), as proof of principle we measured the changes in intracellular metabolites of *E. coli* MG1655 grown in M9 upon stimulation with 100 μM ATP. This showed that the metabolites in treated bacteria were easily discriminated from the unstimulated control (Figure S3A) and revealed shifts in the abundance of several compounds from proteinogenic and non-proteinogenic amino acid, fatty acid, small organic acid, pentose phosphate, nucleotide and nucleoside, cell envelope, co-factor metabolism and redox regulation (Figure S3B-F, Table S2). Intracellular metabolites in bacteria treated with 100 μM or 1 mM eATP were only partially different (Figure S3G-H). Two compounds (cAMP and L-lysine – Figure 3I and Figure S3E, Table S2) were significantly negatively modulated by eATP at both concentrations, while arginosuccinic acid and L-pipecolate decreased at 100 μM eATP (Figure S3E). Interestingly, tyramine accumulated in 100 μM eATP-treated bacteria, as observed for 1 mM eATP, indicating that for some pathways the response is very sensitive. These results show that the response to eATP is concentration-dependent at quantitative level (seen as duration and strength of transcriptional induction) and also leads to cells with different metabolic states.

Altogether, our results show that eATP triggers a strong metabolic response in a non-pathogenic bacterium in a medium- and dose-dependent manner, resulting in the production of bioactive compounds.

### Comprehensive genome-scale modelling reveals medium-specific differences in *E. coli* metabolic capabilities induced by eATP

We integrated transcriptional and intracellular metabolomic data from *E. coli* MG1655 exposed to 1 mM ATP by performing genome-scale metabolic modelling with REMI^60^ to gain a comprehensive understanding of the key properties of eATP-modulated metabolic networks.

The analysis revealed that in LB 78% (141/181) and in M9 59% (138/233) of the omics-based constraints were consistently integrable with the model (Table S3), indicating that nutrient-rich medium provides more metabolic flexibility. We performed differential flux analysis and identified 316 and 430 reaction fluxes that were significantly (*P* value <1e-3) modulated for LB and M9, respectively (Table S3). ATP stimulation enriched the following metabolic subsystems in both LB and M9: valine, leucine and isoleucine metabolism; murein synthesis and recycling; and cell envelope biosynthesis. Glycerophospholipid metabolism was modulated in LB, while in M9 we observed modulation of the core metabolism (pentose phosphate pathway and citric acid cycle), non-essential amino acid metabolism (proline and arginine), purine and pyrimidine nucleotide biosynthesis (Figure 4A-B, Table S3). These analyses further highlight the medium-specific differences in metabolic by eATP and point to bacterial envelope changes, in accordance with transcriptional data (Figure 2J).

**Figure 4.**
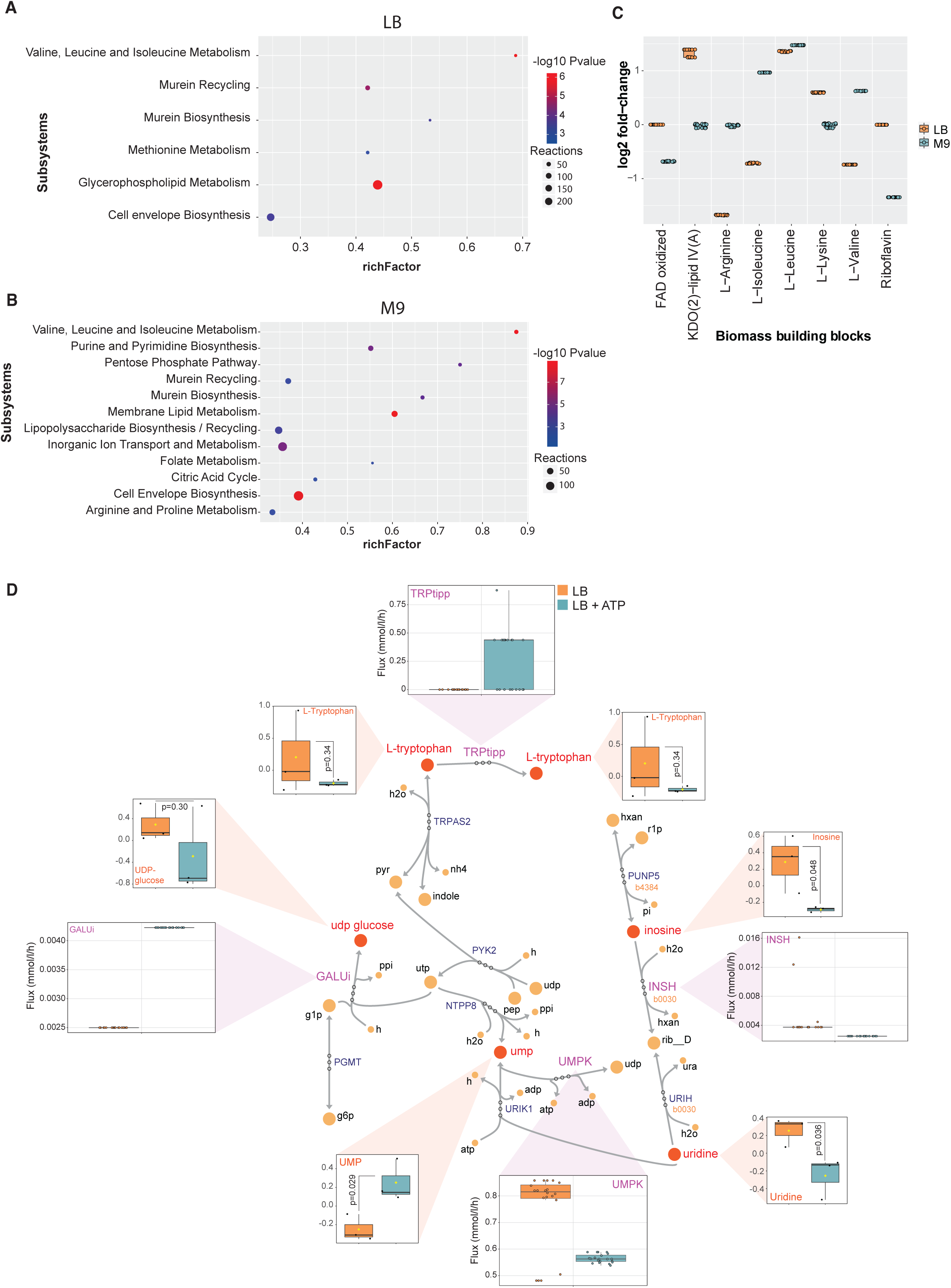
Model-based enrichment analysis of predicted reaction fluxes and analysis of biomass building blocks triggered by eATP in non-pathogenic *E. coli*. **A**, Metabolic subsystems enrichment analysis for *E. coli* MG1655 stimulated with 1 mM eATP for 180 min in LB using differentially regulated reactions (DERs). **B**, Metabolic subsystems enrichment analysis for *E. coli* MG1655 stimulated with 1 mM eATP for 180 min in M9 using differentially regulated reactions (DERs). **C**, Differential fold changes of *in silico* production for eight biomass building block groups between *E. coli* MG1655 grown in LB or M9 in presence of 1 mM eATP for 180 min *versus* the unstimulated control. Each dot is an alternative solution predicted by the genome-scale metabolic model. A cut-off P-value <0,05 was used. **D**, Identification of metabolic subnetworks based on differentially regulated reactions and differentially regulated intracellular metabolites triggered in *E. coli* MG1655 by addition of 1 mM ATP for 180 min in LB (n=3). Shown are the differentially regulated intracellular metabolites (labelled in red, fold change > 1,5, see also Figure 3H), modulated genes (labelled in orange, fold change > 1,5, Table S1) and differentially regulated reactions (labelled in capital purple and black circles, P-value < 1e-3). Other compounds are labelled in orange circles and written in black. Non-modulated reactions are labelled in capital grey and black circles. TRPtipp, L-tryptophan transport out (periplasm); INSH, Inosine hydrolase; UMPK, UMP kinase; GALUi, UTP-glucose-1-phosphate uridylyltransferase (irreversible).

To understand the downstream effects of transcriptional, metabolite and reaction perturbations, we calculated the abundance of 68 Biomass Building Blocks (BBBs), which are important indicators of bacterial physiology, with focus on amino acids, lipids and co-factors (Table S3). eATP caused marked modulation (> 1.5-fold) of 6 BBBs in LB and 5 in M9 (Figure 4C). In both media, the production of L-leucine increased, while L-lysine and KDO(2)-lipid IV(A), an essential component of lipopolysaccharide, were overproduced only in LB. In M9 the production of L-valine and L-isoleucine increased but decreased in LB. Oxidized flavin adenine dinucleotide (FAD) and riboflavin production diminished in M9, as expected from transcriptional (Figure 2I) and metabolomic data (Figure 3F). These findings indicate that BBBs synthesis, and thereby the chemical nature of bacterial structures such as the cell envelope, is altered by eATP in a medium-dependent manner.

Finally, to obtain a metabolic snapshot of the global changes induced by eATP, we identified subnetworks that contain both significantly modulated reaction fluxes and metabolites or genes (Figure 4D for LB and 5 for M9). This very stringent analysis highlights pathways that are consistently altered at many levels. Most reactions were increased upon addition of eATP in both media (seen in iterative quantitative simulations shown in boxes), with a few exceptions: UMPK (see figure legend for abbreviations) and INSH in LB and GLUPRT in M9. eATP modulated the nucleotide salvage network in LB and M9, but the extent and pattern of deregulation differed. Purine-nucleoside phosphorylase-dependent reactions (PUNP2, PUNP5 encoded by *b4384*, and PUNP6 encoded by *b2407*) play a vital role in recycling purines. Expression of *b2407* was not altered by eATP in M9, while *b4384* was downregulated in LB (Table S1). In LB only inosine, UMP, uridine, and few reactions (PUNP5 and INSH) were impacted by eATP. By contrast, in M9 the network showed modular deregulation, with several metabolites (inosine, adenosine, adenine, UMP, uracil) and reactions (PUNP2, PUNP6, DADA, and AMPN) being modulated by eATP. UMP was upregulated in both media, but produced through UPPRT reaction in M9, whilst through NTPP8, UMPK or URIK1 in LB. Moreover, N-acetyl-glutamate, ADP and oxidized glutathione accumulated in M9 in presence of eATP. While there is no direct interaction between N-acetyl-glutamate and oxidized glutathione, they may influence each other indirectly, as the reaction fluxes of GTHS and ACGS were higher in M9 in presence of eATP. Further, D-ribose-5 phosphate increased, together with directly or indirectly linked reactions associated with nucleotide salvage pathways (AMPN), pentose-phosphate pathway (TKT1), co-factor and prosthetic group biosynthesis (NNAM) as well as purine/pyrimidine biosynthesis (GLUPRT). Of note, ATP addition led to a slight but significant decrease of nicotinamide (vitamin B3), a precursor of the essential redox co-factor nicotinamide adenine dinucleotide (NAD).

Taken together, these results highlight the potent impact of eATP on the metabolism of non-pathogenic *E. coli*, including on the production of crucial cellular components and of compounds with known bioactive properties on bacteria or the host. Moreover, these findings underscore the importance of considering the cellular context when analysing changes in metabolic networks induced by eATP.

### eATP alters antibiotics and antimicrobial peptides susceptibility

During infection, bacteria may be exposed to eATP in conjunction with environmental insults such as antibiotic therapy and antimicrobial peptides from the host.

Our results indicated that eATP triggers global physiological changes, some of which trigger envelope stress responses (Figure 2J) and modify the cell envelope (Figure 4A-C) in non-pathogenic *E. coli*. Therefore, we tested the sensitivity of eATP-pretreated *E. coli* MG1655 to selected antibiotics and antimicrobial peptides of diverse origin and modes of action (Table S4). Non-pathogenic *E. coli* showed enhanced susceptibility to aztreonam, mecillinam, and gentamicin and reduced susceptibility to ceftazidime and imipenem (Figure 6A). Bacteria treated with eATP showed a marked increased susceptibility to all tested antimicrobial peptides except CAP-18 (Figure 6B-H and Figure S4A-G).

**Figure 5.**
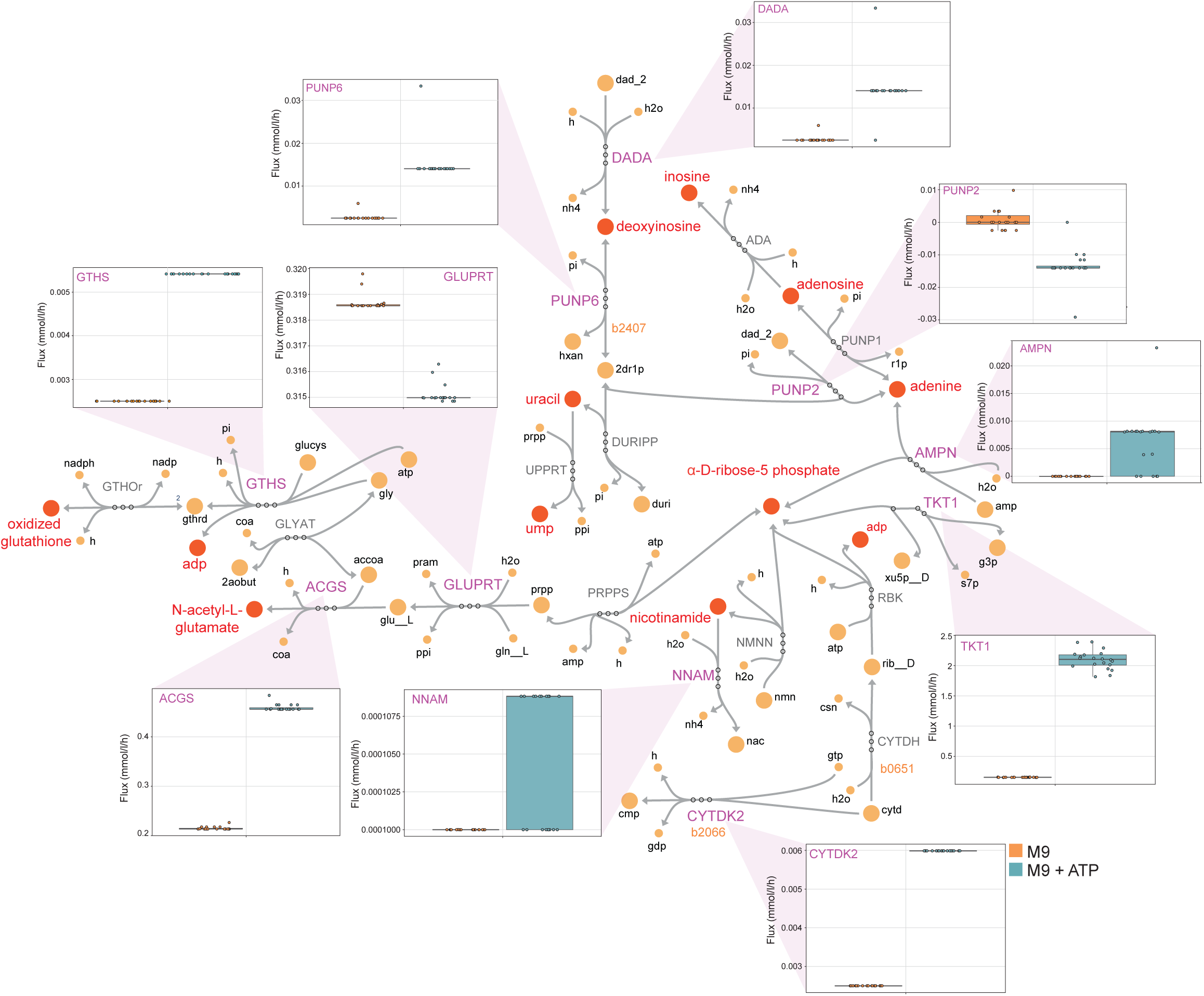
Metabolic subnetwork analysis integrating transcriptional and metabolic changes triggered by eATP in non-pathogenic *E. coli* in M9. Identification of metabolic subnetworks based on differentially regulated reactions and differentially regulated intracellular metabolites triggered in *E. coli* MG1655 by addition of 1 mM ATP for 180 min M9 (n=4). Shown are the differentially regulated intracellular metabolites (labelled in red, fold change > 1,5, see also Figure 3I), modulated genes (labelled in orange, fold change > 1,5, Table S1) and differentially regulated reactions (labelled in capital purple and black circles, P-value < 1e-3). Other compounds are labelled in orange circles and written in black. Non-modulated reactions are labelled in capital grey and black circles. DADA, Deoxyadenosine deaminase; PUNP6, Purine-nucleoside phosphorylase (Deoxyinosine); GLUPRT, Glutamine phosphoribosyldiphosphate amidotransferase; ACGS, N-acetylglutamate synthase; GTHS, Glutathione synthetase; NNAM, Nicotinamidase; CYTDK2, Cytidine kinase (GTP); TKT1, Transketolase; AMPN, AMP nucleosidase; PUNP2, Purine-nucleoside phosphorylase (Deoxyadenosine).

**Figure 6.**
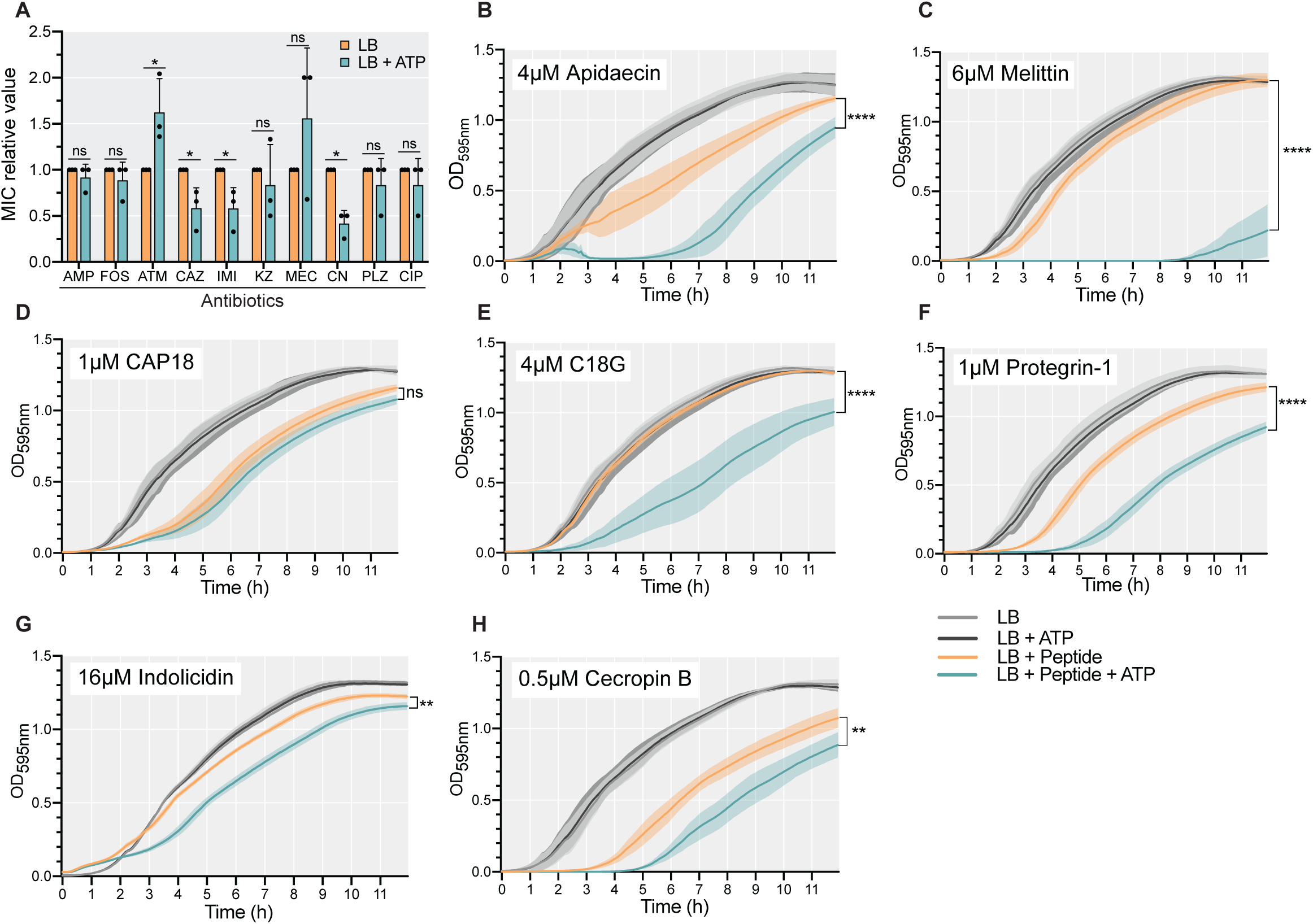
eATP alters antibiotics and antimicrobial peptides susceptibility. **A**, Relative antibiotic sensitivity of *E. coli* MG1655 grown on LB compared to LB + 1 mM ATP plates in the presence of ampicillin (AMP), fosfomycin (FOS), aztreonam (ATM), ceftazidime (CAZ), imipenem (IMI), cefazolin (KZ), mecillinam (MEC), gentamicin (CN), plazomicin (PLZ), or ciprofloxacin (CIP). Shown is the mean with s.d. of 3 independent experiments. Statistical analysis: multiple t-test; *P < 0.05; ns, not significant. **B-H**, Growth of *E. coli* MG1655 in LB upon treatment at time 0 with antimicrobial peptides or vehicle in the presence or absence of 1 mM ATP. Shown is the mean with s.d. of 3 independent experiments performed in duplicates or triplicates. Statistical analysis: two-way ANOVA with Dunnett’s multiple comparison test; **P < 0.01, ***P < 0.001 and ****P < 0.0001; ns, not significant.

Altogether, these data show that, consistently with our metabolic flux and biomass building blocks analyses, in non-pathogenic *E. coli* eATP alters the sensitivity to antibiotics or antimicrobial peptides.

### The ability to respond to eATP is present among clinically relevant bacteria

We explored whether the capacity to respond to eATP by modifying the transcriptional landscape is present in bacteria other than non-pathogenic *E. coli* MG1655. We selected a panel of harmless, pathobiontic or pathogenic Enterobacteriaceae: non-pathogenic *E. coli* BW25113, adherent invasive *E. coli* (AIEC) LF82 (pathobiontic), probiotic *E. coli* Nissle 1917, enteropathogenic *E. coli* (EPEC), enteroinvasive *E. coli* (EIEC), pathogenic *Shigella flexneri* (enteroinvasive), S*almonella enterica* serovar Typhimurium (enteroinvasive) and *Klebsiella pneumoniae*.

As proof of concept, we transformed these organisms with transcriptional fluorescent reporter plasmids carrying promoters from *E. coli* MG1655 that were activated by eATP (Figure 1, Figure S1 and Table S1). To estimate the chances of *E. coli* MG1655 promoters to function in these bacteria, we performed multiple sequence alignments and sequence identity analyses (Figure S5A). This revealed a high percentage of identity among all *E. coli* strains and *S. flexneri*, whereas *S*. Typhimurium and *K. pneumoniae* showed lower identity compared to *E. coli* MG1655. All tested Enterobacteriaceae displayed a transcriptional response to eATP in LB and M9 that was within the same range as *E. coli* MG1655, with a few exceptions (Figure 7A, Figure S5B); P*gadW-gfp* in *E. coli* Nissle 1917 was only activated in M9, while we detected no activation in *S. flexneri* or *S*. Typhimurium. None of the reporters responded to eATP in *K. pneumoniae* grown in M9. We omitted experiments with *S. flexneri*, EIEC and *S*. Typhimurium in M9 owing to their poor growth.

**Figure 7.**
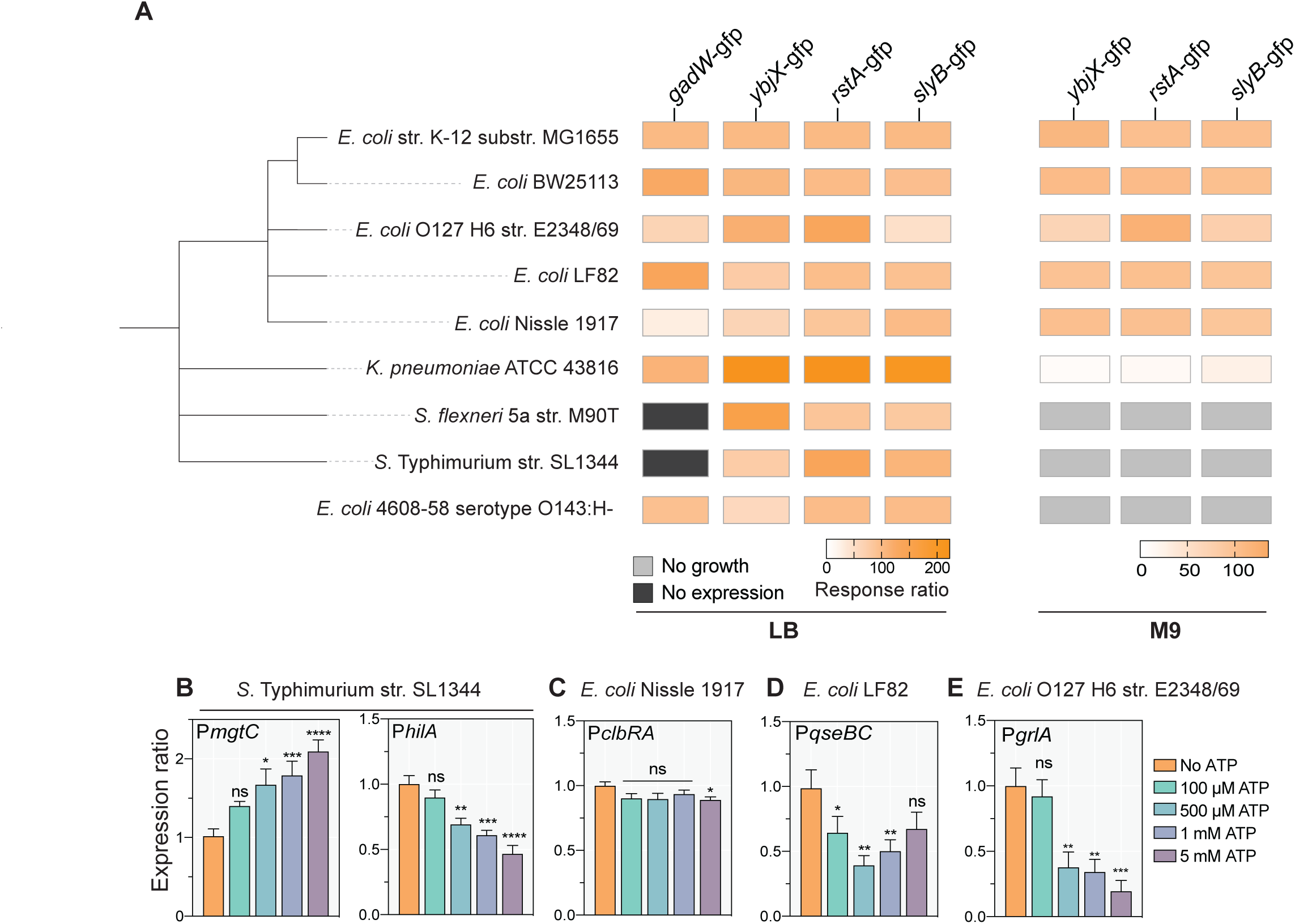
eATP triggers a transcriptional response in several Enterobacteriaceae and regulates the expression of virulence factors. **A**, Response ratio to eATP in percentage in selected bacteria transformed with reporter plasmids encoding promoters that upregulate the expression of GFP in eATP-treated *E. coli* MG1655, which was used as reference (100%). Bacteria were stimulated with 1 mM ATP for 180 min in LB or M9. Shown is the average ratio from ≥ 3 independent experiments performed in triplicates. **B-E** , Promoter activity of selected virulence or fitness factors seen as GFP expression normalised to number of bacteria upon addition of increasing concentrations of ATP for 180 min in LB. Indicated promoter regions were cloned into GFP reporter plasmids and transformed into indicated corresponding strains. Increasing concentrations of eATP were added and the expression ratio was calculated comparing the fluorescence/OD_600nm_ in LB without ATP with fluorescence/OD_600nm_ in LB + ATP. Shown is the mean with s.e.m. of ≥ 3 independent experiments performed in duplicates or triplicates. Statistical analysis: two-way ANOVA with Dunnett’s multiple comparison test; *P < 0.05, **P < 0.01, ***P < 0.001 and ****P < 0.0001; ns, not significant.

These results indicate that eATP is an environmental cue for medically relevant bacteria. While in Enterobacteriaceae the ability to respond to eATP is conserved, the type of response is strain- and species-specific.

### The expression of virulence factors is regulated by eATP

We showed that eATP elicits transcriptional responses in pathogenic Enterobacteriaceae, exploiting target genes that are mostly present in the core genome. To determine whether eATP could also regulate strain-specific virulence genes, we cloned the promoter of selected virulence and fitness factors into the transcriptional reporter plasmid, which we introduced into the bacteria of origin, and monitored promoter activity upon eATP stimulation.

We selected the promoters of *S*. Typhimurium *mgtC*, *hilA*, and *ssrB*. In *Salmonella*, pathogenicity islands 1 and 2 (SPI-1 and SPI-2) have crucial roles in early and later stages of infection, respectively^61^. HilA positively controls the expression of SPI-1 genes^62^, while the two-component system SsrAB regulates the expression of several SPI-2 operons^63^. MgtC aids bacterial survival inside macrophages^64^. Addition of ATP significantly upregulated P*mgtC* and downregulated P*hilA* in a dose-dependent manner (Figure 7B), whilst P*ssrB* was not regulated (Figure S5C).

*E. coli* Nissle 1917 contains a *pks* genomic island that mediates the synthesis of colibactin, a genotoxin^65^. ClbR is a key transcriptional regulator of the *pks* island^66^. Stimulation with high concentrations of eATP resulted in a decrease of the *clbRA* promoter activity (Figure 7C).

We also tested the promoter of *E. coli* LF82 *qseBC*. This strain was isolated from a mucosal lesion of a Crohńs disease patient^67^. It adheres to and invades enterocytes, and replicates within macrophages. The pathogenicity of LF82 and that of many *E. coli* depends on the quorum-sensing two-component system QseBC^68^. Upon treatment with eATP, the promoter activity of LF82 *qseBC* significantly decreased at all eATP concentrations except 5 mM (Figure 7D), in accordance with eATP-dependent regulation of quorum sensing genes in *E. coli* MG1655 (Figure 2H).

Finally, we tested P*ler* and P*grlA* from EPEC E2348/69. EPEC encodes a pathogenicity island, the locus of enterocyte effacement (LEE). LEE is activated by the master regulators Ler and GrlA^69^. While the activity of P*grlA* strongly diminished in a dose-dependent manner (Figure 7E), P*ler* remained unaltered (Figure S5D).

Together, these findings indicate that eATP can regulate virulence and fitness in a broad panel of clinically important intestinal bacteria.

## Discussion

In lower and higher eukaryotes, eATP is a universal signal that regulates their physiology and acts as danger signal^1,2,8^. Here we show that non-pathogenic and pathogenic Enterobacteriaceae are able to respond to eATP. Heightened eATP elicits extensive changes in physiology and virulence by reprogramming transcriptional and metabolic landscapes. The transcriptional response is specific to intact eATP and leads to production of an intracellular second messenger. These effects show that eATP is an environmental cue to bacteria.

We previously discovered that the pathogenic Enterobacteriaceae *Shigella flexneri*, *Salmonella enterica*, and EPEC trigger ATP secretion to the gut lumen as they infect the intestinal epithelium^36,42^. Host cells respond to this danger signal by positively and negatively regulating defence mechanisms. We now show that the very bacteria inducing ATP secretion and non-pathogenic gut bacteria also respond to this host danger signal by up- or downregulating virulence and fitness factors such as biofilms downstream of eATP. These processes are in line with bacterial adaptation to an inflammatory environment.

The dose-dependency of the response indicates that microbes may utilise eATP as a site- and infection stage-characteristic environmental signal, which provides an additional regulatory layer that finetunes species-specific adaptation. Indeed, it’s reasonable to assume that different niches in the gut have distinctive eATP concentrations during inflammation, akin to oxygen tension gradients^70^, as follows. In healthy tissue the extracellular space is virtually devoid of ATP, while injured cells release up to millimolar ATP from their 3-10 mM cytosolic ATP pool^8^. Hence, eATP is likely most abundant at its site of release, the mucosa, and gradually diffuses into the gut lumen. Thus, harmless microbes in the lumen might be exposed to lower eATP than pathobionts or pathogens adhering to the epithelium. Invasive bacteria encounter secreted ATP during cellular uptake^42^ and peaking ATP after vacuolar escape^71,72^. Accordingly, we generally observed downregulation of early-stage and upregulation of late-stage virulence genes, when eATP is highest. In eukaryotes, eATP acts as a signal in the micro-to millimolar range^1–3,8^. Consistent with this and the existence of intestinal eATP gradients, we observed sensitive regulation of metabolic pathways and virulence factor expression at high micro- and millimolar ATP concentration.

Besides regulating virulence and fitness factors, exposure to eATP alters the susceptibility to antibiotics and antimicrobial peptides. This is consistent with remodelling of bacterial structures and building blocks observed by metabolic flux analyses. Of note, elevated eATP augments the production of gut antimicrobial peptides during experimental shigellosis^36^, setting the conditions for the host to exploit enhanced sensitivity as an additional defence mechanism. Importantly, our data indicate that the potency of antimicrobials may be altered in inflammatory environments, where eATP is abundant. This is of therapeutic concern for the antibiotics showing reduced efficacy in the presence of eATP and reinforces the need to test antimicrobials under physiological conditions^73,74^.

eATP-dependent bacterial regulation may take place in various environments. Indeed, eATP is a ubiquitous inflammatory mediator, for example in the airways^75^. *K. pneumoniae* is carried in the respiratory and gastrointestinal tracts^76^. Moreover, heightened eATP in dense microbial communities in or outside the human body could regulate bacterial adaptation, for instance in survivors of bacteriophage- or antibiotic-induced death^18,28,32–34^.

In conclusion, these findings warrant comprehensive investigation of the impact of eATP on diverse microorganisms in various environments. As in eukaryotes, we found that eATP modifies bacterial gene expression. Strikingly, an ATP analogue that binds to mammalian ATP sensors induced transcriptional activation in bacteria. In non-pathogenic *E. coli*, eATP triggered only partially overlapping promoter activities in different media. In fact, the output to an external cue is a global function of distinct input components: the cue and the cellular state. The latter strongly depends on the environment^77^, thus giving rise to media-specific responses. Further, the existence of early and late gene expression programmes reflects the hierarchical structure of transcriptional regulatory systems, which are comprised of environmental stimuli at the top and multiple lower levels of transcriptional regulators and target genes^77^.

Likewise, in different media eATP induced only partially overlapping shifts in compounds of diverse chemical nature and genome-scale metabolic networks. As for transcriptional states, several regulatory mechanisms influence metabolic fluxes, including reactant concentration (depending on the environment and cellular state), enzyme abundance (depending on gene expression), and enzyme kinetic parameters^78,79^, thus explaining medium-specific metabolite changes. Introduction of ATP as a substrate may contribute to the observed changes in intra- and extracellular purines. While eATP is neither transported into *E. coli*^20,55^ nor serves to phosphorylate bacterial molecules^20^, its components may be internalised after ATP hydrolysis. Indeed, carbon flux mapping in intracellular *S.* Typhimurium showed that ribose and nucleobases partly and fully originate from the host, respectively, rather than from *de novo* synthesis^80^. Altogether, our data reveal that the eATP response presents the characteristics of bacterial gene regulatory networks and metabolic fluxes.

Further, eATP stimulated the production of metabolic intermediates or end products with described bioactive properties on microorganisms or the host^81,82^, for example antioxidants and the neurotransmitter tyramine (a complete list is provided in Table S2). Cross-feeding of vitamins is common among intestinal bacteria^69^. Altogether, this points to the activation of intracellular signalling pathways in bacteria downstream of sensing eATP, which culminate in the production of compounds potentially regulating intra- and inter-kingdom cell-to-cell communication.

The mechanism by which bacteria sense and respond to eATP remains to be elucidated. A molecular understanding of bacterial eATP signalling is key to predict which species encode the ability to utilise eATP as environmental cue and to assess, under well-controlled conditions, the role of eATP-mediated host-bacteria interactions in complex systems such as the gut. Our discovery of eATP-dependent transcriptional and metabolic regulation of bacteria, together with its molecular dissection, may open unfathomed opportunities for the prevention and treatment of infection and inflammation.

In conclusion, we show that eATP rewires physiology, antimicrobial sensitivity, and virulence in medically important intestinal bacteria. Thus, eATP emerges as a potent environmental cue in prokaryotes.

## Acknowledgements

A.P. acknowledges institutional support from MIMS (Vetenskapsrådet grant 2021-06602), Umeå University Medical Faculty, Knut and Alice Wallenberg Foundation grant KAW 2015.0225, Umeå University Bioteknikmedel FS 2.1.6-1862.17 and Insamlingsstiftelse FS 2.1.6-452-20, Vetenskapsrådet project grant VR-MH 2022-00778, Carl Kempe Foundation grant SMK-1859, and institutional support from Queen’s University Belfast. S.T. acknowledges Svenska Sällskapet för Medicinsk Forskning (SSMF) for postdoctoral grant P19-0098. We thank David A. Cisneros (Queen’s University Belfast) for critical reading. The graphical abstract was created with BioRender.

## Contributions

Conceptualization: S.T. and A.P.

Data curation: S.T.

Formal analysis: S.T., V.P., M.L.B., M.P.P., C.H.O., A.N.

Funding acquisition: S.T. and A.P.

Investigation: S.T., M.L.B., M.P.P., C.H.O., A.N.

Methodology: V.P.

Project administration: A.P. Resources: O.B.

Software: S.T., V.P. and N.S. Supervision: S.T. and A.P. Validation: S.T. and A.P.

Visualization: S.T.

Writing – Original Draft: S.T., V.P. and A.P.

Writing – Reviewing and editing S.T., V.P., M.L.B., M.P.P., N.S., C.H.O., O.B., A.N., A.P.

## Material and methods Reagents

Unless stated otherwise, all chemicals, antibiotics and primers were purchased from Sigma-Aldrich. Restriction and cloning enzymes were obtained from New England Biolabs (NEB). Plasticware for bacterial culture was purchased from Techno Plastic Products (TPP), Starstedt, and Greiner. ATP sodium salt solution 100 mM was obtained from ThermoFisher (reference R0441). ATPγS (#NU-406-5), BzATP (#NU-1620-5), cAMP (#NU-1503S), cGMP (#NU-1501-50), c-di-AMP (#NU-954S) and c-di-GMP (#NU-951S) were obtained from Jena Bioscience. ADP (#A2754), AMP (#O1930), adenosine (#A9251), inosine (#I4125), hypoxanthine (#H9636), uric acid (#U2625) were purchased from Sigma-Aldrich, and resuspended in purified water just before use. Antimicrobial peptides were purchased from Eurogentec. Antibiotic MIC strips were obtained from Liofilchem.

## Bacterial strains and culture conditions

The bacterial strains used in this study are listed in Supplementary Table 4. Bacteria were preserved as glycerol stocks at -80°C. Bacterial cultures in rich medium were performed in Luria Broth (LB) 1X medium (yeast 5 g/l; tryptone 10 g/l; NaCl 5 g/l). Bacterial cultures in minimal medium were performed in M9 (M9 salts 1X; CaCl_2_ 0,1 mM; MgSO_4_ 2 mM; glucose 0,5 %). Kanamycin (25 µg/ml) was added to the medium when needed. If not specified, overnight bacterial cultures were grown at 37°C in the medium of choice, subcultured 1:100, and grown at 37°C in a shaking incubator at 150 RPM until mid-exponential phase (OD_600nm_=0.3-0.5).

## Screen for genome-wide promoter activity in *E. coli* MG1655 using a transcriptional reporter library

The library of reporter strains, each carrying a low-copy plasmid (pUA66-*mut2GFP*) with a promoter of interest controlling fast-folding GFP (GFPmut2), was previously described^52^ and purchased from Dharmacon. The library was replicated to accommodate positive and negative controls to create a working library, resulting in punctual changes in sample position with respect to the original library. The disposition of samples is provided in Table S1. Reporter strains were inoculated in 96-well plates from frozen stocks and grown overnight in the medium of choice with kanamycin at 37°C in a shaking incubator. Overnight cultures were diluted 1:100 into 150 μl of fresh medium containing kanamycin in black 96-well plates with an optical flat bottom (Greiner). The cultures were incubated at 37°C in a shaking incubator at 150 RPM. When the mid-exponential phase was reached, 50 μl of medium containing 4 mM ATP was added to each well to obtain a final concentration of 1 mM ATP. In each plate, a strain carrying a promoterless plasmid was included as a background control. Optical density (OD_600nm_) and fluorescence intensity (excitation: 485 nm; emission: 535 nm) were measured in an Infinite M200 or a Spark plate reader (Tecan) just before (t=0 min) and after ATP addition (time=5 min) and then measured at intervals of 60 min for 240 min, with incubation at 37°C in a shaking incubator at 150 RPM between measurements. Promoter activities were calculated by subtracting the background fluorescence (promoterless strain) and by dividing the intensity of fluorescence by the corresponding OD_600nm_ as described in^52^. The reproducibility of the screening was verified by plotting the normalised fluorescence of a random plate measured on day 1 against the values on day 2. Promoters showing a final fluorescence value lower than 100 in both ATP-treated and control samples were considered as not expressed. The ratio of the fluorescent signal produced by eATP-treated bacteria and untreated control bacteria was calculated. The screen was performed twice in LB and in M9, hence statistical analysis was not performed.

## Construction of chromosomal reporter fusions

Reporter fusions were constructed using Red recombination as previously described^83^. The kanamycin resistance cassette was amplified from pKD4 with ST1_aph_KpnI_F and ST2_aph_XhoI_R, cloned into a KpnI/XhoI digested pBluescript plasmid and named pBluescript-kan (See Table S5). The same procedure was performed with the PCR-amplified msfGFP Addgene plasmid #29750 and subcloned into EcoRI/HindIII-digested pBluescript-kan. This PCR fragment contained a ribosome binding site upstream of *msfGFP.* The final plasmid construct was verified by sequencing. PCR amplification of msfGFP-Kan was performed using the newly cloned plasmid using ST5_cbpA_sfGFP_aph_F and ST6_cbpA_sfGFP_aph_R as templates. The PCR product was digested with DpnI and transformed into *E. coli* MG1566 carrying pKD56. Colonies were selected on plates containing 50 μg/ml kanamycin and verified by sequencing. The kanamycin resistance gene was removed by FLP-mediated recombination^83,84^.

## Measurement of promoter activity upon addition of ATP or other compounds at small scale

The experimental protocol employed in the transcriptional reporter screen was adjusted to a smaller scale to determine promoter activities upon addition of different concentrations of ATP or of various compounds, in *E. coli* MG1655 carrying chromosomal fusion reporters or in other bacteria. The reporter strains used in these experiments were directly inoculated into 3 ml of appropriate medium with kanamycin (25 μg/ml) for 16 h at 37°C and 150 RPM. The cultures were then diluted to obtain a final OD_600nm_=0,01 using the medium of choice at a final volume of 150 μl per well in black 96-well plates with an optical flat bottom (Greiner) and, as previously described, grown until mid-exponential phase. After incubation, 50 μl of medium containing the appropriate compound concentration was added to each well to obtain the desired concentration in the final volume (200 μl). The optical density at OD_600nm_ and fluorescence at 485 nm excitation and 535 nm emission were measured. Data were analysed as described above (screening section). To determine the specificity of the response to eATP over other compounds, the response to eATP was used as 100% reference. Experiments were performed in at least 3 biological replicates with 2 or 3 technical replicates.

## Measurement of pH in eATP-treated bacteria

To determine whether ATP supplementation modifies the pH of the growth medium, the samples were treated as to quantify fluorescence reporter activation, followed by collection of 2 ml in microcentrifuge tubes. Samples were centrifuged for 3 min at 13000 x *g*. The supernatant was filtered with a 0,22 μm pore size, 13 mm diameter, syringe-driven filter (Millex). A pH meter (VWR pHenomenal) was used to measure the pH. The mean pH of samples with the standard deviation was calculated.

## CRP-regulated genes, Panther and KEGG pathway analyses

To obtain the count of CRP-regulated promoters among eATP-regulated promoters, the list of CRP-regulated genes was retrieved from RegulonDB v12.0^85^ and compared to eATP-regulated promoters (Table S1).

To investigate the biological functions and gene ontology of differentially expressed genes (DEGs, FC > 1,5) upon exposure to 1 mM ATP, DAVID Bioinformatics Resources version 6.8^86^ and the PANTHER classification system version 17.0^87^ were used to functionally annotate and classify genes. The reference list for the GO enrichment analysis contained 1,807 modulated genes of the transcriptional reporter library^52^.

## Quantification of intracellular cAMP by ELISA

The quantification of intracellular cAMP in bacteria was performed as described previously^88^ with minor modifications. *E. coli* MG1655 was grown overnight in LB or M9. The cultures were diluted to OD_600nm_∼0.01 in 3 ml and incubated until mid-exponential growth. If indicated, 1 mM ATP was added, and bacteria were incubated at 37°C under agitation for additional 3 h. After the treatment, bacteria were harvested by centrifugation at 4°C and 13,000 x *g* for 2 min, washed twice with precooled PBS, and resuspended in 500 μl cooled PBS. The bacteria were lysed by sonication on ice (3 cycles of 30 s separated by 45 s pause). The samples were centrifuged for 20 min at 80,000 x *g* at 4°C, the supernatant was collected and used to measure the total protein content and the cAMP concentration. The protein concentration was determined using the Micro BCA^TM^ Protein assay kit (#23235, Thermo Scientific) following the manufacturer’s instructions. The cAMP quantification was performed using the cAMP assay kit (#4339S, Cell Signalling Technology). The absorbance was measured with a Spark microplate reader (Tecan). The protein and cAMP concentrations of experimental samples was extrapolated from standard curves according to the manufactureŕs instructions. The intracellular cAMP concentrations were converted to nmol per mg of protein.

## Quantification of biofilm formation by microtiter plate assay with crystal violet staining

Overnight cultures of *E. coli* MG1655 in LB or M9 were diluted to OD_600nm_∼0.01 using the appropriate medium and incubated until mid-exponential phase at 37°C and 240 rpm. Then, 100 μl of sub-culture were aliquoted into 96-well cell culture plates (Sarstedt). When indicated, 1 mM ATP or mock were added to the medium. The 96-well plates were statically incubated at 25°C for 24 h and biofilms dyed with crystal violet to determine the biofilm biomass. Briefly, planktonic cells were removed by inverting, and surface attached cells were washed twice using deionized water. Each well was supplemented with 120 μl of 1% crystal violet solution and incubated for 15 min. The crystal violet solution was removed by inverting and the wells were washed twice with 200 μl of deionized water. Then, 150 μl of 30% acetic acid solution was added to dissolve the stained biofilms, and the absorbance at 570 nm was measured using a microplate reader (Spark, Tecan) to determine the biofilm biomass.

## Sample preparation for untargeted metabolomics

*E. coli* MG1655 was grown overnight at 37°C in the medium of choice. The cultures were diluted to obtain a final OD_600nm_=0,01 in 20 ml of appropriate medium and incubated until mid-exponential phase at 37°C under agitation. The subculture was divided into 5 ml samples for each condition. ATP was added (100 μM or 1 mM), and the tubes were incubated at 37°C under agitation for 3 h. The time-matched control samples were supplemented with an equal volume of distilled water. Samples of 2 ml were collected in 2 ml microtubes (Sarstedt, ref 72.690.001) 180 min after the addition of the compounds and centrifuged at 13000 x *g* for 2 min. The supernatant of M9-cultured bacteria was filtered (pore size: 0,22 µm, diameter: 13 mm, syringe-driven, Millex) and collected in a sample tube. The bacterial pellets were washed 3 times with ice-cold PBS, keeping cells on ice between centrifugations. Both the supernatant and the pellet were stored at -20°C until analysis. At least 3 biological replicates were prepared for each condition.

## Metabolite extraction for untargeted metabolomics

Bacterial pellets were solubilized using 200 µl 90:10 MeOH:H_2_O and were stored at - 80°C or kept on dry ice. Acid-washed glass beads of 425-600 μm diameter (Sigma Aldrich) were added to samples kept on dry ice to constitute 50% v/v of the 90:10 MeOH:H_2_O cell suspension, and cells were disrupted by shaking at 30 Hz for 2 min (Mixer Mill MM400, Retsch) using prechilled (4°C) holding blocks. Samples were centrifuged at 4°C and 14000 x *g* for 10 min, followed by transfer of the supernatant into an LC‒MS vial (Chromacol, 03-FISV, Thermo Fisher, MA, USA). Samples were evaporated in a centrifugal evaporator (Speed Vac, Savant) at room temperature until dry and stored at -80°C, and reconstituted with 20 µl 60:40 MeOH:H_2_O prior to LC-MS analysis.

## LC‒MS analysis of metabolites

Five microliters were injected into a 1290 Infinity II LC UHPLC system (Agilent, CA, USA) connected to either a 6550 Series or 6540 Series Q-TOF mass spectrometer (Agilent, CA, USA) with a Jet Stream electrospray ionization (ESI) source (Agilent, CA, USA). Data were collected in positive and negative ionization mode with ESI settings: gas temperature 150°C, drying gas flow 16 l/min, nebulizer pressure 35 psi, sheath gas temperature 350°C, sheath gas flow 11 l/min, Vcap 4000 V, nozzle voltage 300 V, Fragmentor voltage 380 V, Skimmer1 45 V and OctapoleRFPeak 750 V. Metabolites were separated using reversed-phase chromatography using an ACQUITY UPLC HSS T3 column with C18 chemistry, 1.8 µM particle size, and 100 Å pore size of 50 mm length and 2.1 mm inner diameter (Waters, MA, USA). For reversed-phase chromatography, elution solvents were (A) H_2_O, 0.1% formic acid and (B) 75:25 acetonitrile:isopropanol, 0.1% formic acid. For separation, the following linear gradient was used (flow rate 0.5 ml/min): min 0: 0.1% B, min 2: 10% B, min 7: 99% B, min 9: 99% B, min 9.3: 0.1% B, min 10.8: 0.1% B. Column re-equilibration occurred from 9.8 to 10.7 min with an increase in flow rate to 0.8 ml/min.

## Metabolite identification and relative quantification

Raw data was processed using Agilent Mass Hunter Profinder software and the metabolites were identified with an inhouse retention time accurate mass library constructed using synthetic standards on the same analytical setup. A retention time window of 0.5 min and a mass accurace tolerance of 5 ppm was employed. A statistics-based analysis of the metabolites was performed using MetaboAnalyst 5.0, a web-based metabolomics processing tool^89,90^. The peak area for each metabolite was uploaded to the MetaboAnalyst program and normalised using log transformed and autoscaling features (mean-centred and divided by the standard deviation of each variable). If values were missing, an estimate was performed using K-nearest neighbours’ imputation. A one-way ANOVA using Tukey’s honestly significant difference test was used to determine significant features. Partial least squares-discriminant analysis (PLS-DA) was used to examine differences between the groups. Metabolites with a *P* value < 0,05 and fold change (FC) > 1,5 were considered significantly modulated. The normalised peak area of each differential metabolite was used to construct the bar plots. Heatmaps were constructed to provide an overview of the metabolome changes, and PLS-DA was used to show the 25 most important features. VIP (Variable Importance in Projection) values (≥ 1.0) and Student’s *t* test (*P* value < 0.05) were combined to identify the differential metabolites. The VIP score is a weighted sum of squares of the PLS loadings. The weights are based on the amount of explained Y-variance in each dimension. Metabolites with high VIP are more important in providing class separation, while those with small VIP provide less contribution.

## Omics data integration into genome-scale models

To investigate the impact of eATP on *E. coli* MG1655 metabolism in depth, a publicly available genome-scale GEM called iML1515^91^ was utilised. Two different growth media, M9 and LB, were separately considered, and the model was adapted accordingly. We created a metabolic model for M9 using minimal growth medium features/nutrition input values, which we referred to as M9GEM. For LB, nutritional input values were assigned from a previous study^92^, and the adapted model was called LBGEM. The REMI method^60^ was used to integrate transcriptional screen data and metabolomics data into M9GEM and LBGEM, allowing for a better understanding of the perturbation effect on metabolism while simultaneously integrating gene expression and metabolomics data. REMI assumes that the reaction fluxes associated with markedly regulated genes are deregulated, and it also integrates intracellular metabolites by constraining reaction fluxes corresponding to differentially regulated metabolites. The main goal of REMI is to ensure the consistency between gene expression, metabolite concentrations, and reaction fluxes. To determine the specific metabolic differences between two conditions (such as eATP and mock-treated control), REMI creates a separate model for each condition and computes a maximum consistency (MCS) score. This score reflects the number of genes/metabolites whose levels are consistent with fluxes, and the constraints can be incorporated into the model. To generate at least 20 alternative sets for the MCS from the given set of constraints, we employed mixed-integer linear programming (MILP). Due to the computational complexity of the optimization process, we selected 20 solutions for statistical analysis.

## Differential reaction flux analysis

To quantify condition-specific changes in reaction fluxes, we obtained a score (R_i_) for each reaction by utilizing the means and variances of flux distributions in eATP-treated and control conditions. To calculate scores for each reaction, we employed the REMI analysis approach^60^, which involves considering alternative solutions of reaction fluxes for two conditions.

To identify markedly regulated reaction fluxes that were significantly altered, we utilized a cut-off of |R_i_|>1. Additionally, we calculated *P* values using a nonparametric test, namely, the Wilcoxon signed rank test. We considered reactions with a *P* value < 1e-3 to be significantly changed.

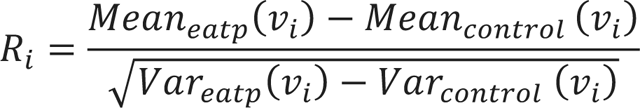

## Pathway and metabolic subsystem enrichment analysis

We conducted pathway enrichment analysis based on modulated genes and metabolites using KEGG pathways as the framework for analysis. To analyse metabolic subsystem enrichment based on metabolic reactions, we utilized GEM iML1515 ^91^. To determine the significance of pathway or metabolic subsystems based on genes, metabolites, or reactions, we employed the hypergeometric probability density function (*P*), as defined by Equation (1):

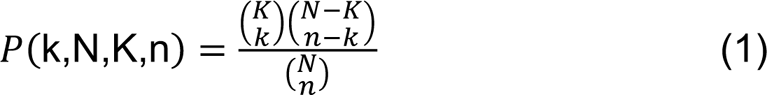

where k and K represent the number of markedly regulated genes, metabolites, or reactions and the total number of genes/metabolites/reactions in a given pathway or metabolic subsystem, respectively. Variables n and N indicate the total number of markedly regulated genes, metabolites, or reactions in the given conditions and the total number of genes/metabolites/reactions in the organism and metabolic context, respectively.

To calculate the *p value* for each pathway or metabolic subsystem, we summed the probabilities for greater than or equal to the enriched number of markedly regulated genes, metabolites, or reactions (as shown in Equation 2):

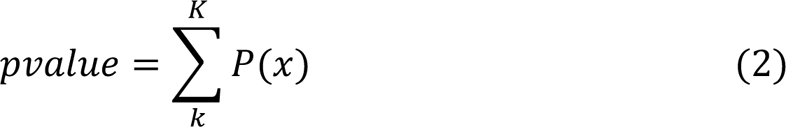

## Biomass building block analysis

Biomass building blocks (BBBs) are crucial components of genome-scale metabolic models that include amino acids, nucleotides, lipids, and other compounds required for cellular growth and maintenance. We used flux balance analysis (FBA), a commonly used constraint-based method, to explore the dynamics of these building components under various environmental conditions. FBA allowed us to examine BBB fluxes in various media, including those subjected to eATP perturbations, providing vital insights into how an organism’s metabolism adapts and responds to these changes.

## Identification of metabolic subnetworks

To gain a more thorough understanding of the metabolic remodelling that results from exposure to eATP, we focused our efforts on identifying subnetworks containing both differentially regulated metabolites (DRMs) and differentially regulated reaction fluxes (DRFs). To accomplish this, we first removed co-factors from the network, as they can be involved in multiple metabolic reactions and can artificially inflate connectivity. This allowed us to better identify important metabolic subnetworks and reveal the true structure of the network. Additionally, we examined the first and second neighbour reactions of each DRM. First neighbour reactions are those that are directly involved with a metabolite’s first neighbours, while second neighbour reactions involve the metabolite’s second neighbours, which are not directly connected to the metabolite but are connected to its first neighbours. Finally, we analysed the most perturbed or deregulated reactions to identify the metabolic subnetworks where the most deregulation occurs in terms of both metabolites and reactions.

## Susceptibility tests to antibiotics and antimicrobial peptides

*E. coli* MG1655 was grown overnight at 37°C in LB. The cultures were diluted to obtain a final OD_600nm_=0,01 in 10 ml of medium and incubated until mid-exponential phase at 37°C under agitation. The subculture was divided into 3 ml samples for each condition. ATP (1 mM) or mock were added, and the tubes were incubated at 37°C under agitation for 3 h.

Resistance to various antibiotics was determined on agar using MIC Test Strips (Liofilchem). Exponential cultures were adjusted to OD_600nm_=0,1 in LB and homogeneously spread onto an LB agar plate containing or not 1 mM ATP. The strip was applied once the plates had fully dried. Plates were incubated over night at 37°C and the MIC at which the no-growth halo was observed was determined visually.

Susceptibility to antimicrobial peptides (Eurogentech) was tested by adjusting the OD_600nm_=0,04 in LB, 50 μl of culture were distributed into a 96-well plate (TPP), followed by 150 μl containing serially diluted antimicrobial peptides together with vehicle or 1 mM ATP. Plates were maintained at 37°C with 3 mm linear agitation in a Spark multimode plate reader (Tecan) and the OD_600nm_ was acquired every 10 min. Seven antimicrobial peptides were used: indolicidin, apidaecin IB, CAP-18, cecropin, melittin, protegrin-1, C18G (see also Table S4). The antimicrobial peptide stock solutions were prepared beforehand in sterile water and kept at −80°C until usage, except for protegrin-1 and C18G, which were prepared in 0,01% acetic acid before freezing. Experiments were performed in biological triplicates with 2 or 3 technical replicates.

## Cloning of promoter regions into pUA66-*GFPmut2*

The high-fidelity polymerase Phusion was used for all PCRs. All plasmid constructs were verified by sequencing (Eurofins). Plasmids and oligos are listed in Table S5. Promoter regions (positions with respect to start ATG) of *mgtC* (-480 to 113), *grlA* (- 89 to 143), *hilA* (-700 to 457), *clbRA* (-537 to 156), *qseBC* (-480 to 109), *ler* (-407 to 108) and *ssrAB* (-431 to 112) were PCR amplified with primers containing either *XhoI* or *BamHI* restriction sites, using DNA extracted from the relevant organisms with the Wizard® genomic purification kit (Promega) according to manufactureŕs instructions. The purified PCR products were XhoI/BamHI digested. The plasmid pUA66-mut2GFP was XhoI/BamHI digested and purified in a 1% agarose gel run at 100 V for 40 min in TAE buffer, followed by isolation with the GeneJet gel extraction kit (ThermoScientific). The digested PCR products and backbone plasmid were ligated using T4 DNA ligase (Thermo Fisher Scientific). The mixture was transformed into CaCl_2_-competent *E. coli* DH5α. Transformants were selected on LB agar plates containing 25 μg/ml kanamycin. A few colonies were picked and screened by colony PCR using primers (Table S5) designed for the upstream and downstream regions of the two restriction sites. Verified recombinant plasmids were purified and transformed into the organisms of promoter origin by electroporation.

## Generation of a phylogenetic tree and percentage identity calculation

A phylogenetic tree of the employed strains based on NCBI taxonomy was generated with PhyloT (https://phylot.biobyte.de/index.cgi). The visualization of the tree was achieved using iTOL (https://itol.embl.de/)^93^. Taxonomy numbers used for the tree are shown in Table S5. To calculate the percentage identity of *gadW*, *ybjX*, *slyB* and *rstA* between the different strains, the nucleotide sequences were downloaded for every strain and aligned using MUSCLE v5 (https://www.ebi.ac.uk/Tools/msa/muscle/). The identity of each sequence compared to the selected reference sequence (*E. coli* MG1655) was calculated using The Sequence Manipulation Suite (https://www.bioinformatics.org/sms2/ident_sim.html) ^94^.

## Statistical analysis

Except for the metabolomic analysis, which was performed using Metaboanalyst (5.0)^90^, all statistical analyses were performed in GraphPad Prism (version 9.0.2). The type of analysis and test of significance is mentioned in the respective figure legends. The default *P* value system of GraphPad Prism was used in all cases and is indicated in figure legends.

## Source code availability

The script for metabolic modelling can be found at https://gitlab.com/obilab1/externalatpmets

## Softwares used

The metabolomic analysis as performed with MetaboAnalyst 5.0^89^. For the visualisation of enrichment pathways and BBB analysis, Python (3.9) and R (4.2) were used. For modelling simulation, IBM CPLEX 12.9 MILP solver and MATLAB r2019a (The Mathworks, Natick, MA, USA) were used. Figures 4D and 5 were created with Adobe Illustrator.

---

**Figure S1.**
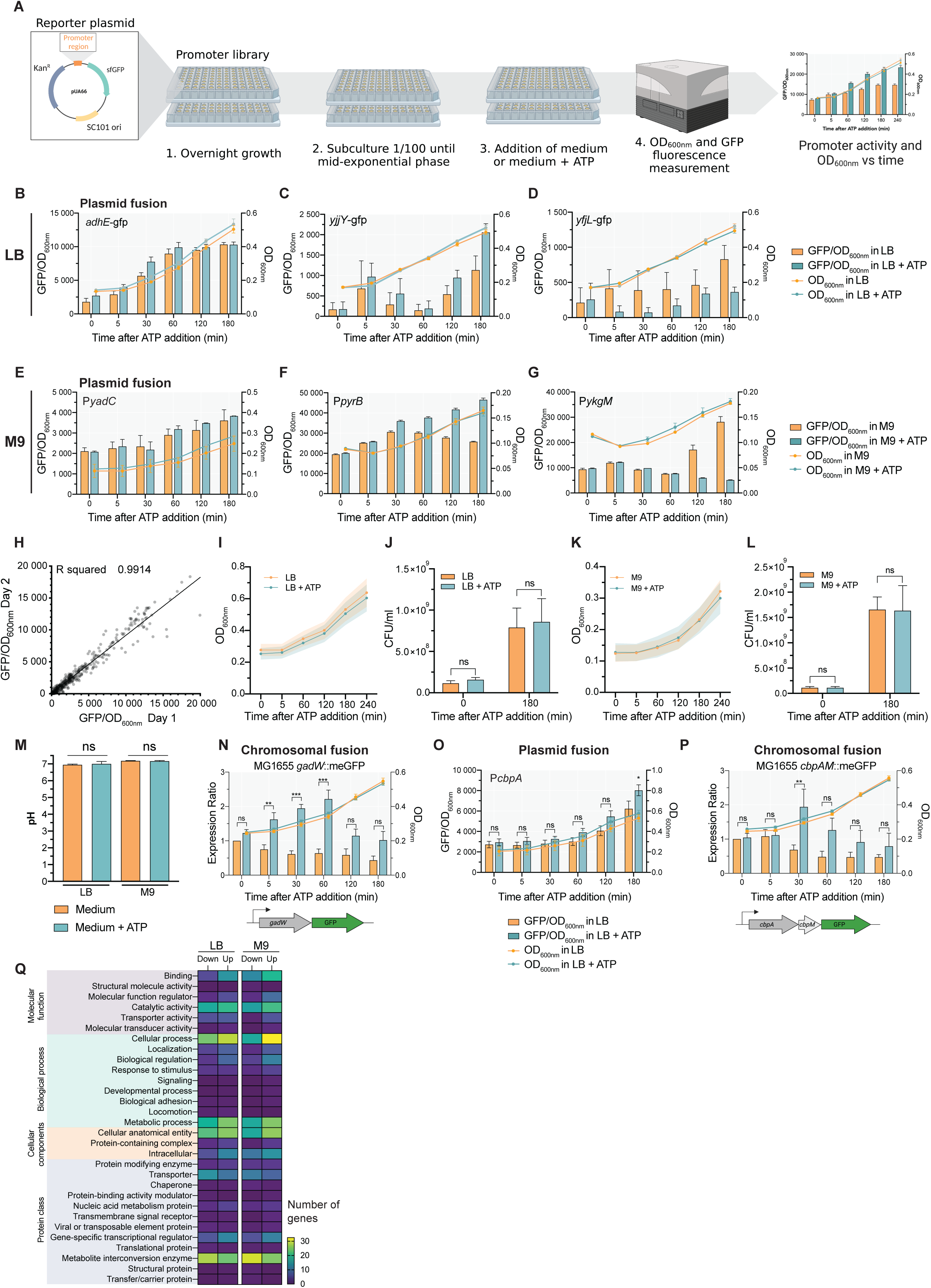
eATP triggers a strong transcriptional response in non-pathogenic *E. coli*, related to Figure 1 and Figure 2. **A**, Overview of the protocol for the promoter library screening for a transcriptional response to eATP. Created using BioRender. **B-G**, Activity of indicated promoters controlling the expression of the reporter GFP (plasmid construct) in the presence or absence of 1 mM ATP obtained in LB (upper panels) or M9 (lower panels). Shown is the ratio of GFP fluorescence to number of bacteria determined by absorbance at 600 nm over time (columns representing the mean with s.e.m., left Y axes), as well as the bacterial growth (continuous lines representing means with s.e.m., right Y axes). **H**, Day-to-day reproducibility for a set of 96 random samples. Shown is the GFP fluorescence divided by the number of bacteria determined by absorbance at 600 nm in LB measured on day 1 and day 2. **I**, Growth curves of *E. coli* MG1655 over 240 min with (blue line) or without (orange line) 1 mM ATP in LB obtained for a set of 2 random plates from the library. The continuous lines represent the mean absorbance and the colour-matched shaded areas the s.e.m.. **J**, Bacterial counts determined by plating before and 180 min after 1 mM ATP addition to *E. coli* MG1655 in LB. **K**, Same as in (**I**) but in M9. **L**, Same as in (**J**) but in M9. **M**, pH of *E. coli* MG1655 cultures after 3 h of incubation with vehicle or 1 mM ATP. **N,** Promoter activity of MG1655 carrying the P*gadW*-mEGFP chromosomal fusion in the presence or absence of 1 mM ATP obtained as in **B-G**. The mean fluorescence value of the unstimulated control at 0 min was set to 1 to calculate relative changes of the fluorescence. The activity of the *gadW* promoter obtained using the reporter plasmid is shown in Figure 1I. **O**, Promoter activity (GFP/OD_600nm_) of the reporter plasmid encoding P*cbpA*-*gfp* in the presence of 1 mM ATP in *E. coli* MG1655. **P**, Promoter activity of *E. coli* MG1655 carrying the P*cbpA*-mEGFP chromosomal fusion in the presence or absence of 1 mM ATP obtained as in **B-G**. The mean fluorescence value of the unstimulated control at 0 min was set to 1 to calculate relative changes of the fluorescence. The activity of the *cbpA* promoter obtained using the reporter plasmid is shown in (O) for comparison. **Q**, Gene ontology analysis using the PANTHER classification system of up- or downregulated genes in LB and M9. Processes with a P value < 1x10^-3^ are shown. Statistical analysis in **N-P**: two-way ANOVA with Dunnett’s multiple comparison test; *P < 0.05, **P < 0.01 and ***P < 0.001; ns, not significant.

**Figure S2.**
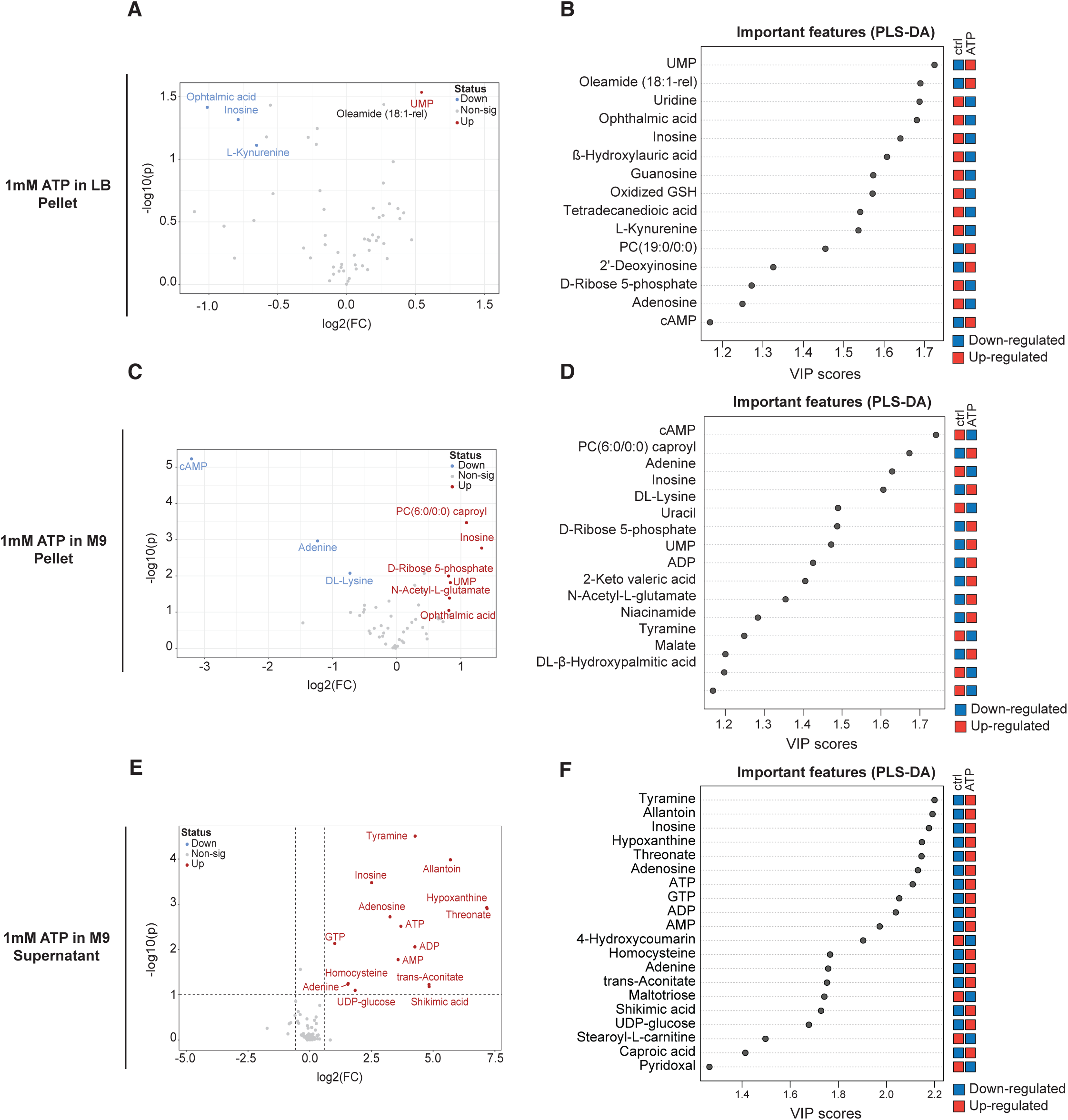
eATP triggers a strong metabolic response in non-pathogenic *E. coli*, related to Figure 3. **A**, Volcano plot analysis of intracellular metabolites changes after addition of 1 mM ATP in LB. Fold change (FC) ≥ 1.5 or ≤ 0.5 (p ≤ 0.05) is considered significant. **B**, VIP (Variable Importance in Projection) score of PLS-DA analysis from samples in (**A**). **C**, Volcano plot analysis of intracellular metabolites changes after addition of 1 mM ATP in M9. Fold change (FC) ≥ 1.5 or ≤ 0.5 (p ≤ 0.05) is considered significant. **D**, VIP score of PLS-DA analysis from samples in (C). **E**, Volcano plot analysis of extracellular metabolites changes after addition of 1 mM ATP in M9. Fold change (FC) ≥ 1.5 or ≤ 0.5 (p ≤ 0.05) is considered significant. **F**, VIP score of PLS-DA analysis from samples in (**E**).

**Figure S3.**
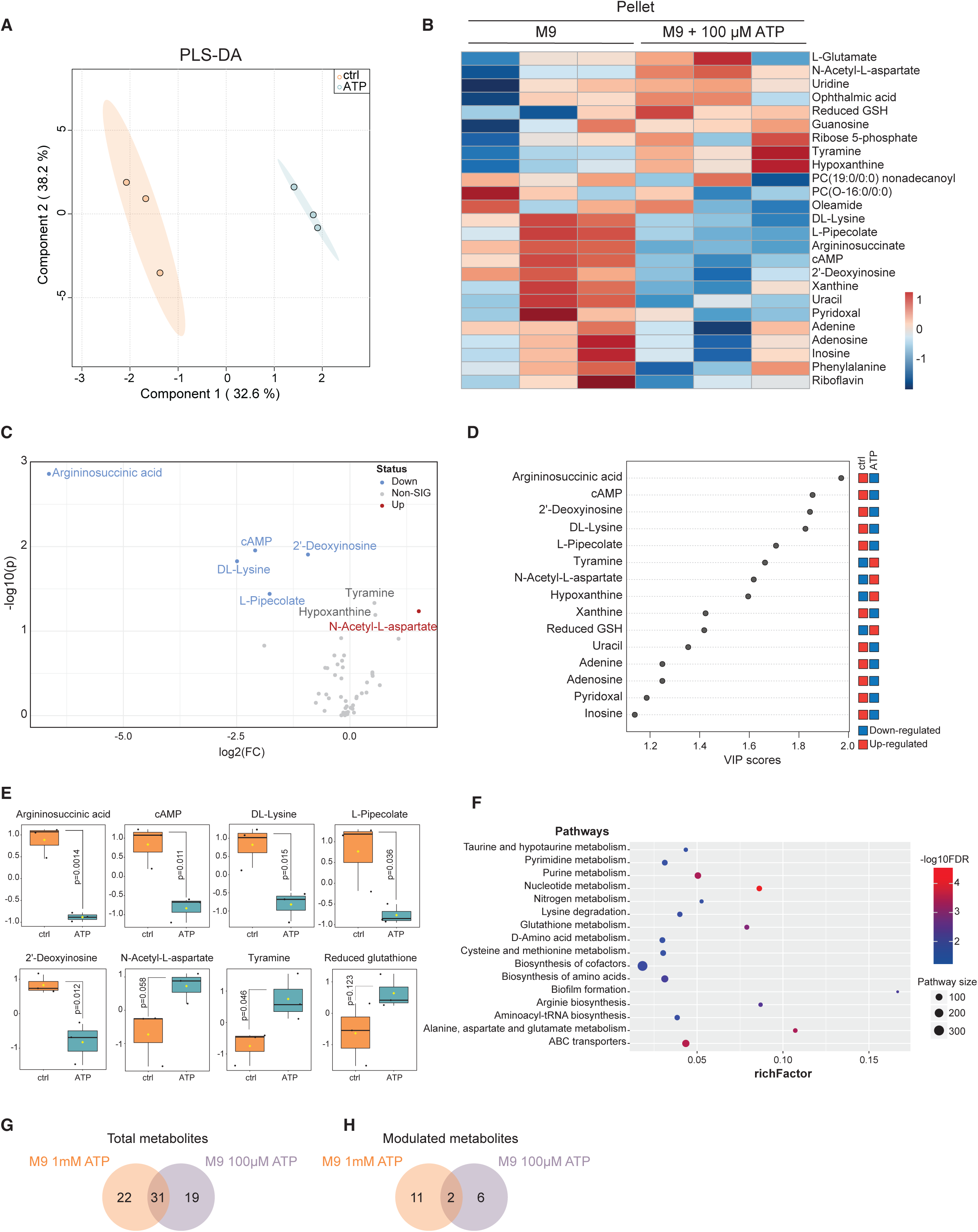
Modulated intracellular metabolites upon exposure to 100 μM ATP in M9 in non-pathogenic *E. coli*, related to Figure 3. **A**, PLS-DA scores plots of components one and two, comparing metabolomic samples (pellet) from treatment with 100 μM ATP for 180 min versus the unstimulated controls (n=3). **B**, Comparison of fold changes in abundance of top 25 metabolites based on PLS-DA analysis from samples in (**A**). **C**, Volcano plot analysis of intracellular metabolites changes from samples in (**A**). Fold change (FC) ≥ 1.5 or ≤ 0.5 (p ≤ 0.05) is considered significant. **D**, VIP (Variable Importance in Projection) score of PLS-DA analysis from samples in (**A**). **E**, Boxplots of relative concentrations of selected modulated intracellular metabolites from samples in (**A**). The bar plots show the normalised values (mean with s.d.). The Y-axis represents normalised peak area of metabolites. Black dots represent the normalised concentrations of all samples of each metabolite. The notch indicates the 95% confidence interval around the median. The mean concentration of each group is shown by a yellow diamond. Statistical analysis: two-sided Welch’s t test; raw P-values are and indicated in each panel. **F**, KEGG pathway enrichment analysis of the up- and down-regulated intracellular metabolites in M9 180 min after 100 μM ATP addition in comparison to unstimulated bacteria. **G**, Comparison of the number of all measured intracellular metabolites in *E. coli* MG1655 180 min after addition of 1 mM or 100 µM ATP M9 medium. **H**, Comparison of the number of all modulated intracellular metabolites in *E. coli* MG1655 180 min after addition of 1 mM or 100 µM ATP M9 medium.

**Figure S4.**
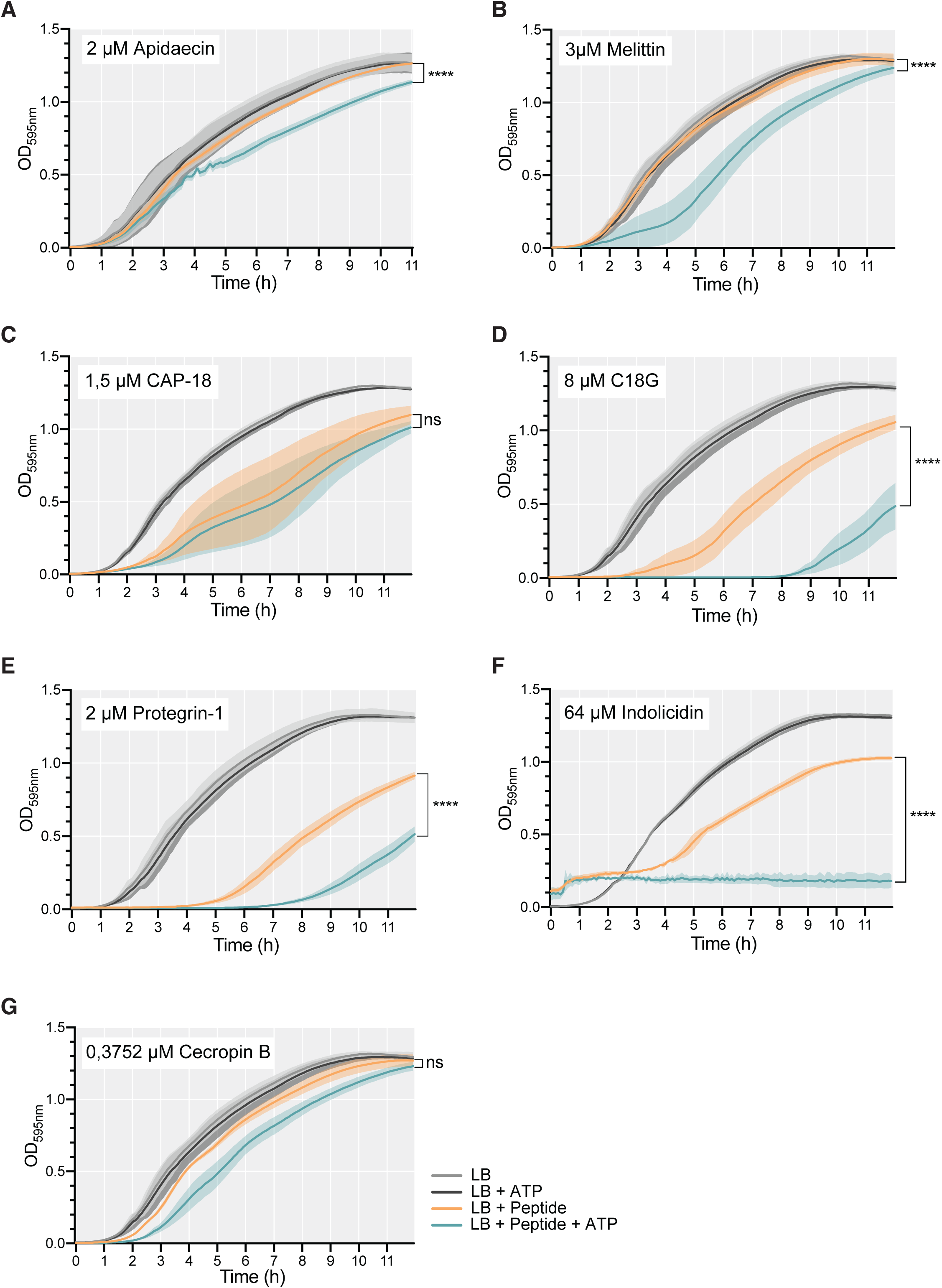
eATP enhances antimicrobial peptide susceptibility, related to Figure 6. **A-G**, Growth of *E. coli* MG1655 in LB upon treatment at time 0 with antimicrobial peptides or vehicle in the presence or absence of 1 mM ATP. Shown is the mean with s.d. of 3 independent experiments performed in duplicates or triplicates. Statistical analysis: two-way ANOVA with Dunnett’s multiple comparison test; ****P < 0.0001; ns, not significant.

**Figure S5.**
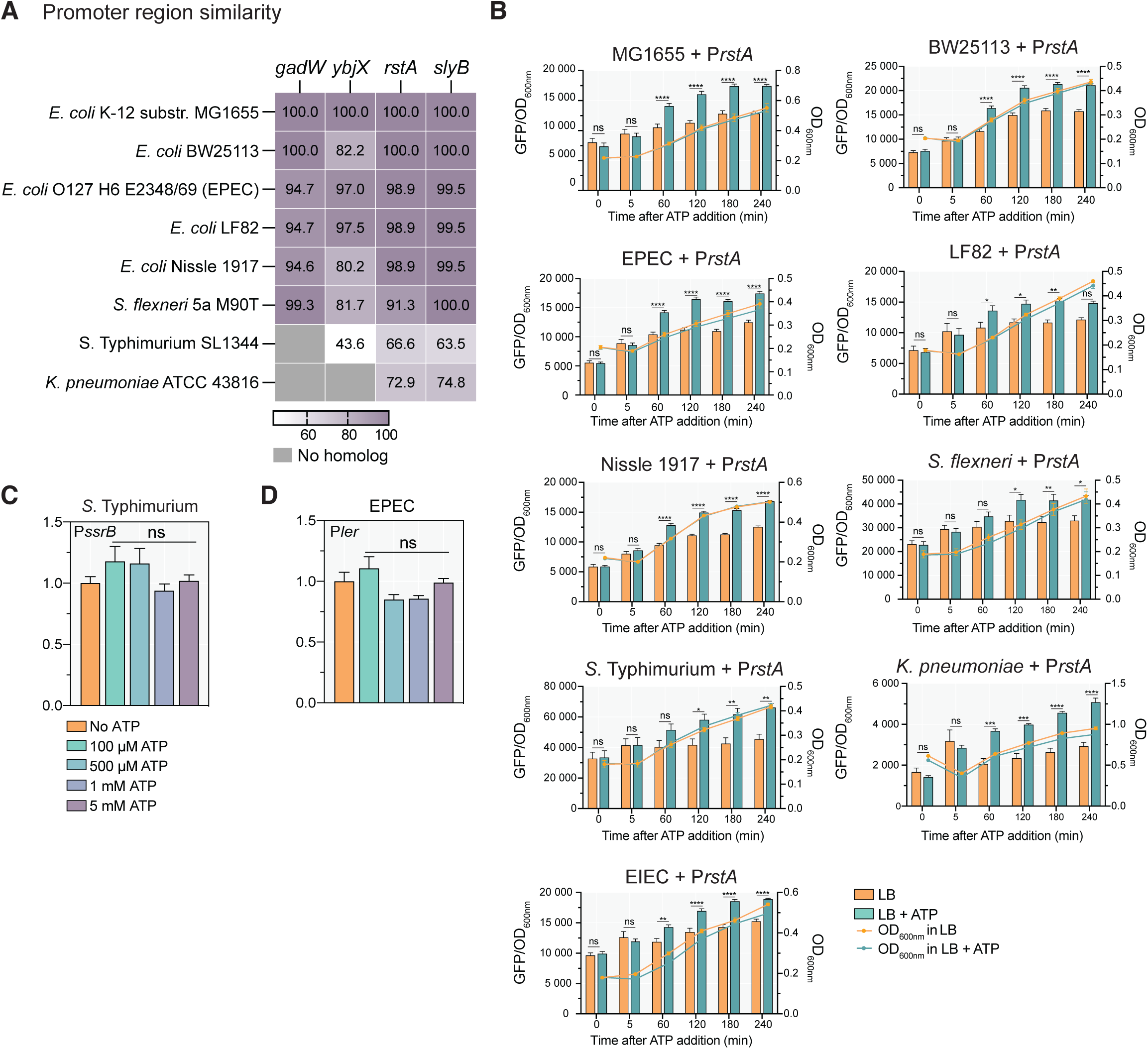
eATP triggers a transcriptional response in non-pathogenic, pathobiontic, or pathogenic bacteria and regulates the expression of virulence factors, related to Figure 7. **A**, Sequence similarity in percentage of promoters P*gadW*, P*ybjX*, P*rstA* and P*slyB* promoters in the indicated bacteria compared to *E. coli* MG1655. **B**, Example of promoter activity over time in indicated strains of using P*rstA* controlling the expression of the reporter GFP in the presence and absence of 1 mM ATP in indicated strains grown in LB, as in Figure 7A. Shown is the ratio of GFP fluorescence to number of bacteria determined by the absorbance at 600 nm over time (columns representing the mean with s.e.m. of 3 independent experiments performed in triplicates, left Y axes), as well as the bacterial growth (continuous lines representing means with s.e.m., right Y axes). Statistical analysis: two-way ANOVA with Dunnett’s multiple comparison test; *P < 0.05, **P < 0.01, ***P < 0.001 and ****P < 0.0001; ns, not significant. **C-D**, Promoter activity of selected virulence factors seen as GFP expression normalised to number of bacteria upon addition of increasing concentrations of ATP for 180 min in LB. The expression ratio was calculated comparing the fluorescence/OD_600nm_ in LB without ATP with fluorescence/OD_600nm_ in LB + ATP. Shown is the mean with s.e.m. of ≥ 3 independent experiments performed in duplicates or triplicates. Statistical analysis: two-way ANOVA with Dunnett’s multiple comparison test; ns, not significant.

**Table S1: Transcriptional reporter library screening** Screening_LB: Summary of the data from the transcriptional library screening in LB. Shown is the average GFP fluorescence divided by OD_600nm_ of at least 2 biological replicates for each time point (0, 5, 30, 60, 120, and 180 min after ATP addition), the fluorescence ratio between the eATP-treated and the mock-treated samples and the log_2_ of this ratio.

Screening_M9: Same as aforementioned but in M9. Panther analysis: Results from the Panther analysis in LB and M9 of differentially expressed genes (DEGs, FC > 1,5) upon exposure to 1 mM ATP for 180 min. LB_genes_KEGG: Pathway enrichment analysis based on differentially expressed genes upon exposure to 1 mM ATP in LB for 180 min. M9_genes_KEGG: Pathway enrichment analysis based on differentially expressed genes upon exposure to 1 mM ATP in M9 for 180 min.

**Table S2: Metabolomic data** Metabolites_LC_MS: Summary of all metabolites identified in *E. coli* MG1655 upon exposure to 100 μM or 1 mM ATP for 180 min and their associated FC and P-value compared to mock-treated bacteria. Metabolites_role: Summary of modulated bioactive metabolites and their roles.

**Table S3:** Metabolic and flux modelling

M9_diff_flux: List of reaction fluxes identified in *E. coli* MG1655 upon addition of 1 mM ATP for 180 min in M9. M9_flux_enrichment: Metabolic subsystems enrichment analysis for *E. coli* MG1655 stimulated with 1 mM ATP for 180 min in M9 using differentially regulated reactions (DERs). M9_BBB: List of the tested biomass building blocks (BBBs) and their associated production rate ratio in eATP-treated compared to mock-treated *E. coli* MG1655 in M9 upon exposure to 1 mM eATP for 180 min. LB_diff_flux List of reaction fluxes identified in *E. coli* MG1655 upon addition of 1 mM ATP for 180 min in M9. LB_flux_enrichment: Metabolic subsystems enrichment analysis for *E. coli* MG1655 stimulated with 1 mM ATP for 180 min in LB using differentially regulated reactions (DERs). LB_BBB: List of the tested biomass building blocks and their associated production rate ratio between the eATP treated *E. coli* MG1655 and the mock in LB upon exposure to 1 mM eATP for 180 min.

**Table S4: List of antibiotics and antimicrobial peptides used in the study**

Antibiotics: List and targets of antibiotics used in the study and the MIC obtained on LB agar and LB agar supplemented with 1 mM ATP. AMPs: List of antimicrobial peptides used in the study.

**Table S5:** Strains, plasmids, and primers

Strains: List of bacterial strains used in the study. Plasmids: List of plasmids used in the study. Primers: List of the primers used in the study.

